# Configurational entropy is an intrinsic driver of tissue structural heterogeneity

**DOI:** 10.1101/2023.07.01.546933

**Authors:** Vasudha Srivastava, Jennifer L. Hu, James C. Garbe, Boris Veytsman, Sundus F. Shalabi, David Yllanes, Matt Thomson, Mark A. LaBarge, Greg Huber, Zev J. Gartner

## Abstract

Tissues comprise ordered arrangements of cells that can be surprisingly disordered in their details. How the properties of single cells and their microenvironment contribute to the balance between order and disorder at the tissue-scale remains poorly understood. Here, we address this question using the self-organization of human mammary organoids as a model. We find that organoids behave like a dynamic structural ensemble at the steady state. We apply a maximum entropy formalism to derive the ensemble distribution from three measurable parameters – the degeneracy of structural states, interfacial energy, and tissue activity (the energy associated with positional fluctuations). We link these parameters with the molecular and microenvironmental factors that control them to precisely engineer the ensemble across multiple conditions. Our analysis reveals that the entropy associated with structural degeneracy sets a theoretical limit to tissue order and provides new insight for tissue engineering, development, and our understanding of disease progression.

## INTRODUCTION

Tissue structure, defined as the number, composition, and spatial arrangement of cells, has reproducible features (order) yet remains heterogeneous in its details (disorder). Individual cells, for example, can differ significantly in their pattern of contacts with other cell types, their shape, and their polarity despite having reproducible average properties. These and other types of spatial heterogeneity are well appreciated in the breast^1^, pancreas^2^, liver^3^ and intestine^4^, and temporal heterogeneity occurs during development^5,6^ and in response to stimuli^7–9^. Order in the spatial arrangement of cells is critical for tissue function^10,11^, and therefore, it may not be surprising that disorder can become accentuated in disease states like cancer^12–16^. Despite the ubiquity of structural heterogeneity in health and disease, its fundamental drivers remain poorly understood.

Heterogeneity in the local arrangement of cells within tissues can be a consequence of extrinsic factors like microenvironmental variability and intrinsic factors like the stochasticity of cellular processes^17,18^. A systematic analysis of these extrinsic and intrinsic factors is limited by the complexity of the microenvironment and the challenges associated with measuring and perturbing live tissues in vivo. In vitro systems such as organoids – self-organizing tissues that closely mimic the structure and function of their source organ^19^ – offer a powerful alternative. Many extrinsic factors like cellular composition, tissue geometry, organization of the extracellular matrix (ECM), and the concentration of diffusible factors can be controlled in organoids using modern bioengineering techniques^20–22^. Remarkably, however, the resulting structures remain heterogeneous^23–26^, pointing to the importance of intrinsic factors including stochasticity in the position and timing of cell proliferation and differentiation^1,27,28^, among others.

Structural order at the tissue-scale emerges from a combination of the mechanical and biochemical properties of single cells and their interactions, which together contribute to their programs of self-organization^29^. How these mechanisms drive the emergence of order at the tissue-scale has been the subject of extensive investigation both experimentally and theoretically, but the opposing cell- and tissue-intrinsic mechanisms promoting disorder have been explored less thoroughly. For example, mathematical models of cell sorting successfully predict the gross structural motifs of tissue when parameterized only with the energy associate with cellular contacts^30,31^, but fail to quantitatively reproduce the average arrangement of cells or its variance. Computational implementation of these models provides a means of incorporating stochasticity or random noise as a way to navigate metastable states during self-organization^27^, but the cellular basis of this noise and its quantitative mapping to tissue-scale order and disorder has not been established. Therefore, it remains unknown precisely how patterns of order and disorder at the tissue scale emerge from the measurable properties of single cells. Consequently, we are unable to understand or engineer these fundamental features of tissues effectively.

To establish a quantitative link between the heterogeneity in tissue structure and the properties of single cells, the mammary epithelium provides an ideal model system. The structure, cell compositions, signaling, mechanics and dynamics of the mammary gland are well characterized both in vivo and in vitro^31–35^. Moreover, the broad rules through which the branched structure of the organ forms^1^, as well as the detailed mechanics through which cell positioning is regulated^31^, are well understood. Mammary organoids recapitulate multiple features of glandular biology in vivo and can be manipulated genetically and with a variety of bioengineering techniques^32^. Like other organoid systems, variability across a number of structural metrics spontaneously emerges in reconstituted mammary organoids even when starting from similar initial conditions^31,36^, suggesting that structural heterogeneity can emerge intrinsically during their self-organization.

Here, we deploy a variety of bioengineering techniques to limit the impact of extrinsic sources of tissue structural heterogeneity on reconstituted human mammary organoids by controlling their initial size, composition, and microenvironment. This allows us to quantify the processes promoting order (tissue surface energies) and opposing processes promoting disorder (stochastic cellular dynamics and entropy). We use a maximum entropy formalism to derive the quantitative relationship between the biophysical and biomolecular properties of single cells and the probability distributions of tissue structure, thereby enabling the systematic engineering of the structural ensemble. Our analysis explains why observed tissue structures can deviate significantly from those predicted by energy-based models of tissue self-organization^30,37,38^. More broadly, it reveals that the capacity of self-organization to maintain order at the tissue-scale is fundamentally limited by fluctuations at the cellular-scale. We anticipate these conceptual and mathematical tools will find applications as diverse as regenerative medicine, tissue modeling, and disease prevention.

## RESULTS

### Cell positioning is intrinsically heterogeneous in vivo and in vitro

Structural order and disorder can be measured using a variety of quantitative metrics^1,39,40^. A metric central to the ability of the mammary gland to synthesize and pump milk is its bilaminar arrangement of luminal and myoepithelial cells (LEP and MEP, respectively)^41^, which can be quantified by the proportion of LEP in contact with the basement membrane (**Fig. 1A**, **Supp. Fig. 1A**, **Supplemental Information**). This structural motif is shared by numerous other secretory organs including the prostate^42^, salivary^43^ and lacrimal glands^44^, and occurs frequently during development. Analysis of normal human breast tissue sections revealed the mean LEP contact with the basement membrane (LEP boundary occupancy or ф_b_) differed significantly from tissue composition (LEP proportion or Φ), indicating an active role for self-organization in promoting structural order (**Fig. 1A**, images from Shalabi et al.^45^). However, it also revealed high variability in ϕ_b_ (**Fig. 1A**). The variability can be partially ascribed to local differences in cell proportions, geometry, signaling and microenvironment^33,46^, which are challenging to constrain in vivo. Surprisingly, however, we noted that regions with similar composition and geometry remained heterogeneous (**Supp. Fig. 1B**, for e.g., mean, 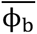 = 0.19 and standard deviation, σ = 0.19 for regions with Φ ≈ 0.5), suggesting additional sources of variability remain.

**Figure 1:**
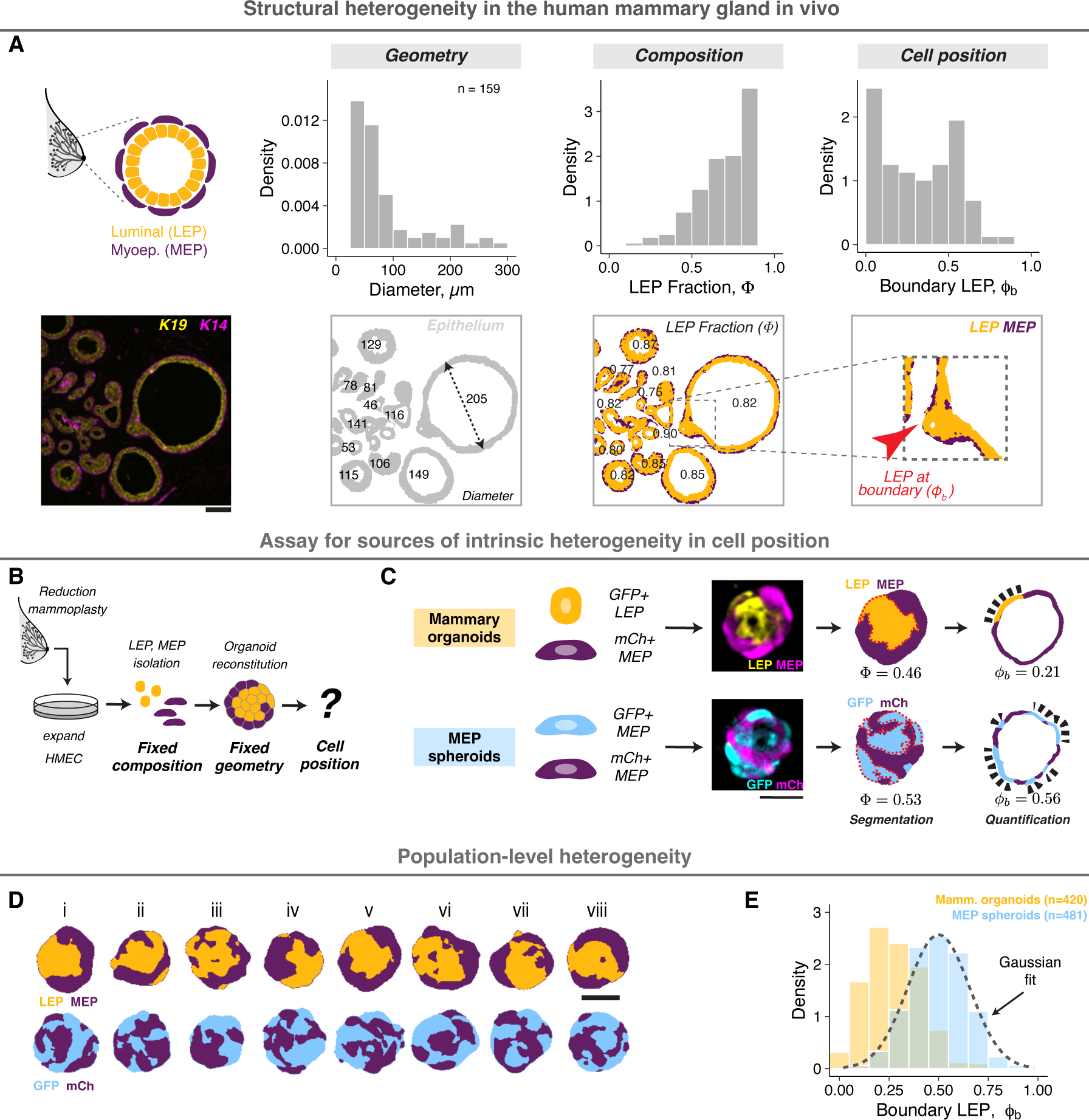
Cell positioning is intrinsically heterogeneous in vivo and in vitro. A. A representative section of normal human mammary gland stained for keratin 19 and 14 (LEP and MEP markers, respectively). The cells are arranged in a bilaminar structure with MEP surrounding LEP (order), however the tissue also exhibits large variance in local geometry, cell proportions and cell positioning (disorder). A custom analysis workflow was used for pixel segmentation and image quantification (**Supplemental Information**). The density histograms show the distributions for effective tissue diameter (d), LEP proportion (Φ) and LEP positioning at the tissue boundary (ϕ_b_). Analysis of n=128 tissue objects from 14 donors is shown. Scale bar = 50 µm. B. Reconstituted organoids provide an in vitro model to study intrinsic sources of positional heterogeneity in tissues with defined composition and geometry. Finite-passage human mammary epithelial cells (HMEC) were isolated from breast reduction mammoplasty, expanded in vitro, sorted as single cells and reaggregated in defined numbers and proportions. Organoids were cultured in Matrigel for 2 days. C. Mammary organoids contained similar number of GFP+ LEP (gold) and mCh+ MEP (purple). MEP spheroids contained similar number of GFP+ MEP (blue) and mCh+ MEP. Confocal images were processed to quantify the total LEP/GFP fraction (Φ) and the boundary LEP/GFP fraction (ϕ_b_) (**Supplemental Information**). For each organoid, three central sections spaced 5 µm apart were analyzed. Scale bar = 50 µm. D. Processed mammary organoid and MEP spheroid images following segmentation illustrate population-level structural heterogeneity. Scale bar = 50 µm. E. Probability density histograms showing the population distribution of mammary organoids (gold) and MEP spheroids (blue) two days post-assembly. The dashed line represents a Gaussian fit to the MEP spheroid distribution. The number of observations is noted at the top right of the graph.

To further constrain extrinsic sources of variability we used primary human mammary ^31^organoids where cell state is stable on experimental timescales^31,47^, and organoid geometry, cell proportion and microenvironment can be independently specified using bioengineering techniques (**Fig. 1C**). Human mammary epithelial cells (HMEC) were isolated from breast reduction surgeries and expanded through several passages. We aggregated equal numbers of LEP and MEP (tagged with GFP and mCherry, respectively) in microwells, then transferred the organoids to laminin-rich hydrogels (Matrigel) for culture and imaging (**Supp. Fig. 1C,D**). These organoids self-organize under the influence of interfacial cellular mechanics to exclude LEP from the tissue boundary within two days^31^, without any significant changes to cell state (**Supp. Fig. 1E**). The boundary occupancy of LEP (ϕ_b_) (**Fig. 1D, Supplemental Information**) was directly analogous to the structural metric we measured in vivo and correctly distinguished well-organized from poorly-organized organoids (**Supp. Fig. 1F-J**). While individual organoids were heterogenous in their detailed structure (**Fig. 1E,F**), they had a reproducible average (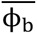 = 0.27, σ = 0.14) over time (**Supp. Fig. 1J**), across cells from different donors (**Supp. Fig. 1K**), and over experimental replicates (**Supp. Fig. 1L**). Additionally, the average in vitro structure was similar to the average in vivo structure for regions with similar size and composition (**Fig. 1B**). Thus, live imaging of reconstituted organoids enables the more systematic analysis of intrinsic sources of variability acting during mammary epithelial self-organization.

The difference between the boundary and total proportion of LEP in mammary epithelial organoids was consistent with the notion that active processes promote order during their self-organization. However, it also raised the question of how we might quantify order and disorder in this system on an absolute scale. To answer this question, we began by investigating the form of a maximally disordered distribution of organoid structures, which we generated by preparing spheroids containing equal numbers of GFP- and mCherry-tagged MEP (**Fig. 1C**). In these MEP-only spheroids, all cells were mechanically and molecularly identical, and therefore, all possible cellular arrangements were equally probable. The average ϕ_b_ of MEP spheroids now matched the cellular composition (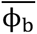 = 0.5, σ = 0.14, **Fig. 1D,E**) and the distribution had a Gaussian form, the maximum entropy distribution for a given mean and variance. This apparent discrepancy between the observed (Gaussian) and expected (uniform) distributions implied the existence of multiple degenerate cellular configurations (also known as **microstates**), all having the same measured ϕ_b_ (defined as a **macrostate**). Macrostates comprising higher numbers of individual microstates would be both statistically and entropically favored^48,49^. This reasoning suggested that the principle of entropy maximization could represent a powerful paradigm for modeling organoid structural distributions.

### Tissues dynamically sample from the ensemble steady state distribution

Cells and tissues exist far from thermodynamic equilibrium, for which the principle of entropy maximization is typically valid^50^. However, the steady state of a non-equilibrium system can be modeled using equilibrium statistical mechanics given certain assumptions hold at the relevant time and length scales^51–54^. These assumptions include the time invariance of distributions and averages as well as the absence of flux across macrostates (also known as “balance”). To investigate whether these assumptions hold for mammary organoids at the steady state (day 2), we first measured temporal variation in the organoid structure across minutes to hours where the confounding effects of cell proliferation and death were minimal (**Supplemental Information**). The structure of both mammary organoids and MEP spheroids fluctuated around distinct averages that did not change over time (**Fig. 2A, Supp. Fig. 2A**). The time-averaged distribution of each was qualitatively similar to the steady state distribution of the larger ensemble, suggestive of the ergodic-like behaviors characteristic of equilibrium systems (**Fig. 2B**). The organoids also had minimal net flux between structural macrostates, as indicated by a diagonally symmetric transition probability matrix at short- and long-time intervals (20 minutes and 3 hours, respectively) (**Fig. 2C, Supp. Fig. 2B**). Additionally, organoids that transiently deviated away from the average structure relaxed back towards it within 3-5 hours (**Fig. 2D**), with relaxation times comparable to the decay time of the autocorrelation function for ϕ_b_ (**Supp. Fig. 2C**). Finally, the steady state distribution was independent of the starting structures, as pre-segregated MEP spheroids made by combining smaller GFP+ and mCh+ cell aggregates also relaxed to a similar distribution as those starting as random structures (**Supp. Fig. 2D**). These combined analyses confirmed the existence of a steady state in these organoids, supporting the applicability of an equilibrium statistical mechanics framework to model the heterogeneity of organoid structural ensembles.

**Figure 2:**
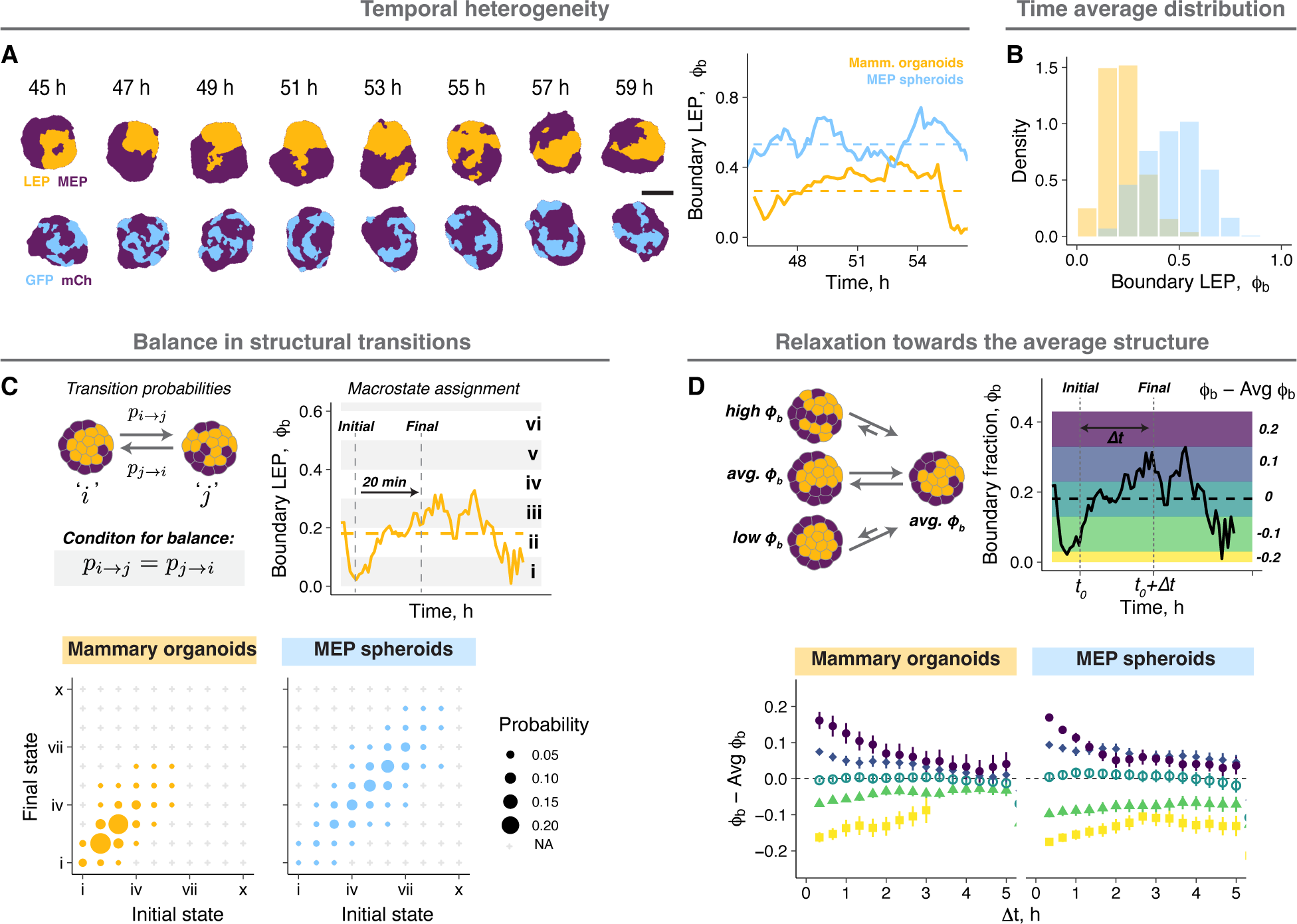
Tissues dynamically sample from the ensemble steady state distribution. A. Snapshots from time lapse microscopy after segmentation for a representative mammary organoid and MEP spheroid illustrate temporal structural heterogeneity at the steady state. Scale bar = 50 µm. Quantification of fluctuations in ϕ_b_ over time for the examples shown. Dashed line is the average of ϕ_b_ over time for the corresponding tissue. B. Probability density histograms showing the temporal distribution of a small number of mammary organoids (n=18) and MEP spheroids (n=24) at the steady state. C. Organoids at different timepoints were binned into 10 structural states according to their ϕ_b_. The probability of transitioning between any two structural states over a 20 min window is represented by the size of the circles. Any transitions not observed during this window are marked by ‘+’. For organoids at the steady state, the diagonal symmetry of the transition probability matrix suggests there is no net flux across states. The same tissues were used for this analysis as Fig. 2B. D. Organoids at each time point were classified into 5 groups based on the difference between the instantaneous and average ϕ_b_. For each bin, the average structure at different time intervals from the initial classification is plotted. The colors in the graph on the right represent the bins on the example trace. The same tissues were used for this analysis as panel Fig. 2B.

### A statistical mechanical framework provides a quantitative description of organoid structural distributions

The principle of maximum entropy allows statistical inference of the most likely distribution of microscopic states given partial information about the system (e.g., an average energy or concentration)^48,49^. In Boltzmann statistics, this distribution corresponds to one that maximizes entropy for a fixed average energy among particles, and the probability of different states depends on their energy and temperature. Analogously, the maximum entropy distribution for a given average organoid structure also resembles a Boltzmann distribution, but where the average energy of the tissues is predicted to be proportional to ϕ_b_ and with a minimum corresponding to ϕ_b_ = 0 (**Supplemental Information**). We therefore asked whether we could identify and measure properties of mammary organoids that functioned analogously to an average energy, the distribution of microscopic states, and temperature in Boltzmann statistics.

Tissue interfacial mechanics are the driving forces of cell sorting^30,37,38^, the process through which mammary organoids and many other tissues^10,31,55,56^ self-organize. During cell sorting, organoids trend towards structures that minimize the total surface energy across all cellular interfaces^38^. Accordingly, we hypothesized that the same surface energies also determine the average tissue mechanical energy, and therefore the probability, of different structures. The mammary organoids have two types of cell-ECM and three types of cell-cell interfaces, each with its unique interfacial tension (γ) (**Fig. 3A**). We calculated all five tensions using the Young’s equation, micropipette aspiration and contact angle analysis (**Supplemental Information**, **Fig. 3B-E**). Remarkably, both computational (**Supplemental Information**, **Fig. 3F,G, Supp. Fig. 3A-E**) and mathematical (**Supplemental Information**) analyses revealed that the average macrostate energies were proportional to ϕ_b_ and had a minimum at ϕ_b_ = 0, precisely as predicted by entropy maximization. The energies for different microstates within a macrostate were similar and symmetrically distributed around their average energy (**Fig. 3G, Supp. Fig. 3D**), allowing us to use a mean field approximation to estimate the average macrostate energy^57^. We defined the slope of the average energy with respect to ϕ_b_ as the ***mechanical potential* Δ**E, which was roughly proportional to the difference between γ_LEP-ECM_ and γ_MEP-ECM_ (**Fig. 3G, Supplemental Information**).

**Figure 3:**
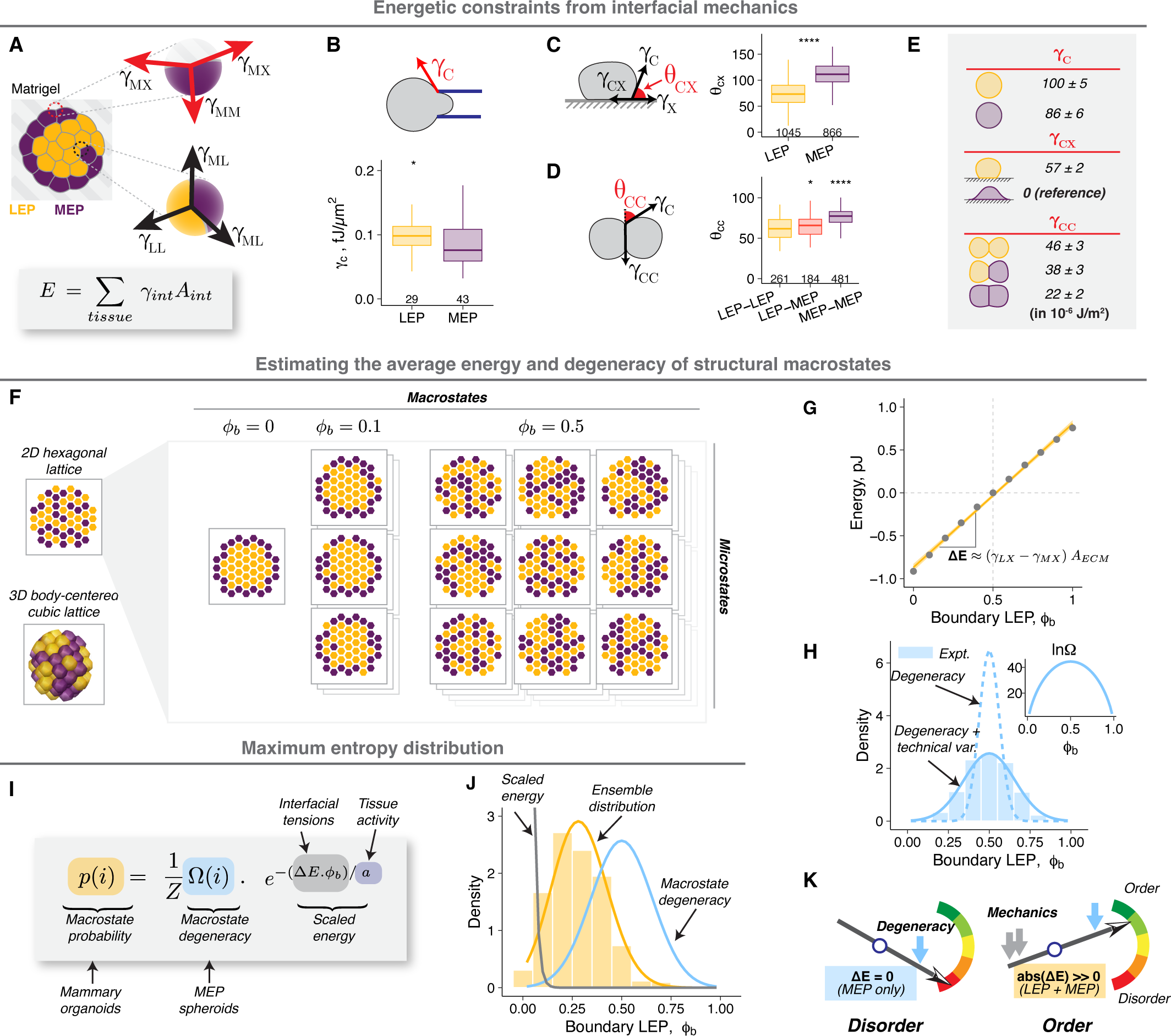
A statistical mechanical framework provides a quantitative description of organoid structural distributions. A. Schematic illustrating tensions at different cell-cell and cell-ECM interfaces. The total tissue mechanical energy is the sum of interfacial energy at each interface (product of the tension, γ_int_, and the area, A_int_, of the interface). B. Cortical tensions of single LEP and MEP in suspension as measured by micropipette aspiration. C. Cell-ECM contact angles for cells on Matrigel-coated glass were measured after 4 h. D. Cell-cell contact angles for cell pairs were measured after 3 h. E. Estimated cell-cell and cell-ECM tensions for LEP and MEP based on Young’s equation. For cell-ECM tensions, the γ_MEP-ECM_ was used as the reference and was assigned the value of 0. Confidence intervals were calculated using error propagation for standard error on cortical tension and contact angle measurements (**Supplemental Information**). F. 2D hexagonal or 3D body-centered cubic (BCC) lattice models were used to estimate the average mechanical energy and the degeneracy of structural macrostates (ϕ_b_) (**Supplemental Information**). Only the 2D model is shown here for simplicity. Macrostates with ϕ_b_ ≈ 0.5 comprise the greatest number of microstates (highest degeneracy). G. The average mechanical energy of mammary organoids for different values of ϕ_b_ estimated from the BCC model. Ten thousand tissue configurations were sampled for each ϕ_b_. The dots and error bars represent the mean and standard deviation. The gold line represents a linear fit for average macrostate energy against ϕ_b_. The slope (ΔE) is roughly proportional to the product of the difference in cell-ECM tensions and the total ECM surface area. H. Macrostate degeneracy (Ω) was calculated analytically (inset) (**Supplemental Information**). The corresponding probability density assuming random sampling of all microstates is shown with the dotted line. Additional variance due to uncertainty in measurements and degeneracy along other structural metrics was built into the model (**Supplemental Information**), and its prediction is shown using the solid line. The superimposed histogram for comparison is the measured ensemble distribution of MEP spheroids (from Fig. 1E). I. The structural distribution of organoid ensembles is modeled as a maximum entropy distribution, a function of the macrostate degeneracy (calculated analytically or from the distribution for MEP spheroids), mechanical energy (calculated from interfacial tensions), and tissue activity. J. The maximum entropy model (gold line) was fit to the measured ensemble distribution of mammary organoids (histogram, from Fig. 1E) to estimate the tissue activity. The predictions for distributions arising from only the scaled energy or macrostate degeneracy are also shown for comparison (gray and blue lines respectively). K. The diagram illustrates how the relative weights of the mechanical energy and macrostate degeneracy determine the extent of structural order. In the absence of a mechanical potential, the degeneracy dominates, and the system is maximally disordered. A large absolute mechanical potential drives the ensemble to an ordered state. The lines and hinges for boxplots in panels **B-D** show the median and the 1^st^ and 3^rd^ quartiles. The number of observations for panels **B-D** are noted at the bottom of the graphs. Asterisks represent the significance of difference from the reference group (MEP for **B** and **C**, MEP-MEP for **D**), as follows ns: p > 0.05; *: p < 0.05, **: p < 0.005; ***: p < 0.0005 based on Wilcoxon test.

In systems at the steady state, microstates with the lowest energy are the most likely to be observed. Macrostates, however, comprise multiple microstates, and their probability is additionally dependent on their degeneracy (Ω, number of microstates). Because MEP spheroids lack any controlling mechanical potential, we reasoned that their observed probability distribution (**Fig. 1E**) is an exact measure of the relative degeneracy of macrostates in mammary organoids. To test this notion, we applied a similar analytical and computational approach as with the energy calculations (**Supplemental Information, Fig. 3F,H, Supp. Fig. 3F-L)**. We predicted minimum and maximum values of Ω at ϕ_b_= 0 or 1 and ϕ_b_= 0.5 respectively, matching the experimental distribution of MEP spheroids (**Fig. 3H**). The strong agreement between predictions and experiments suggests the model fully captures the degeneracy of the configurational phase space with no additional hidden degrees of freedom. Importantly, we confirmed that Ω is only a function of cell proportions and tissue geometry and not cell identity (**Supp. Fig. 3M**). Therefore, this approach allows a priori estimation of macrostate degeneracy and the maximum entropy distribution in the absence of any mechanical driving forces, in contrast to inferring entropy from measured distributions^48^.

The calculated ΔE and Ω can be used to estimate the macrostate probabilities by modeling the ensemble structural distribution as a maximum entropy distribution, similar to a Boltzmann distribution (**Fig. 3I, Supplemental Information**), but requires we define an additional scaling factor for the mechanical energies analogous to temperature in equilibrium thermodynamics. At higher temperature, higher energy states are increasingly occupied, thereby decreasing order in the system. Temperature derives from the average kinetic energy of the microscopic parts of the system, and manifests as fluctuations across configurations with different energies. In the context of non-equilibrium biological systems like tissues, the magnitude of these fluctuations is instead primarily determined by the active mechanics of cellular interfaces^58^, which are impacted by stochastic processes associated with the actomyosin dynamics that power cell motility, contractility, division, apoptosis, and endocytosis^59–62^. Therefore, we defined ***tissue activity*** as an energy scaling factor, which we estimated by fitting organoid distributions to a maximum entropy distribution parameterized with the calculations for ΔE and Ω (**Fig. 3I,J, Supp. Fig. 3N**). We note that the inferred activity (∼ 10^-13^ J) is many orders of magnitude higher than the energy scale of molecular thermal fluctuations (k_b_T ∼ 10^-21^ J at 300 K), but of a similar scale to measured interfacial energies (**Fig. 3E**), and therefore consistent with active processes. The magnitude of mechanical energies relative to activity determine the energetic constraints for the structural ensemble, regulating the extent of structural order (**Fig. 3K**).

### Tissue activity sets the balance between the mechanical potential and macrostate degeneracy

We next aimed to directly test whether activity is a tunable physical property of cells that sets the balance between Δ**E** and Ω. We reasoned that cell motility would be a measurable parameter associated with activity, as activity should scale with the average kinetic energy, or the integrated motion of cells within organoids (**Fig. 4A**). MEP consistently had a higher average speed than LEP, suggesting they were the primary source of tissue activity. Moreover, MEP proximal to the tissue boundary moved the fastest (**Fig. 4A**), an observation consistent with other studies showing higher cell motility at tissue surfaces^63–65^. Therefore, we hypothesized that the primary driving forces of MEP motility, and thus tissue activity, are tractions between MEP and the ECM. Consistent with this hypothesis, cell motility was greatly reduced in organoids cultured in non-fouling agarose microwells as assessed by calculating an effective diffusion coefficient of cell nuclei (∼2-fold reduction) (**Supplemental Information, Fig. 4B-C, Supp. Fig. 4A,B**). To test the impact of reduced activity on ensemble structure, we examined mammary organoid self-organization in agarose microwells (**Fig. 4D**). Under these conditions we predicted two important impacts on steady state tissue organization: reduced activity due to the loss of ECM tractions but also an altered mechanical potential due to the loss of cell-ECM adhesion, which is a principle driving force for self-organization. We first confirmed that the measured macrostate degeneracy does not change under these conditions (**Supp. Fig. 4C,D**). We also confirmed that ΔE is significantly reduced in magnitude and has a negative slope in agarose (**Fig. 4E**), which without a corresponding reduction in activity, would result in only weak sorting of LEP to the tissue boundary. Surprisingly, however, we observed a strong boundary enrichment of LEP (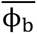 = 0.82, σ = 0.11, **Fig. 4F, Supp. Fig 4C**). The unexpectedly high LEP enrichment implied a 5-fold lower activity in agarose compared to Matrigel (**Fig. 4F**). The lower predicted activity was consistent with reduced structural fluctuations (**Fig. 4G, Supp. Fig. 4E**) and the lower diffusion constant in agarose (**Fig. 4C**). Therefore, an increase in structural order can also emerge in tissues with small mechanical driving forces, so long as tissue activity is sufficiently reduced (**Fig. 4H**).

**Figure 4:**
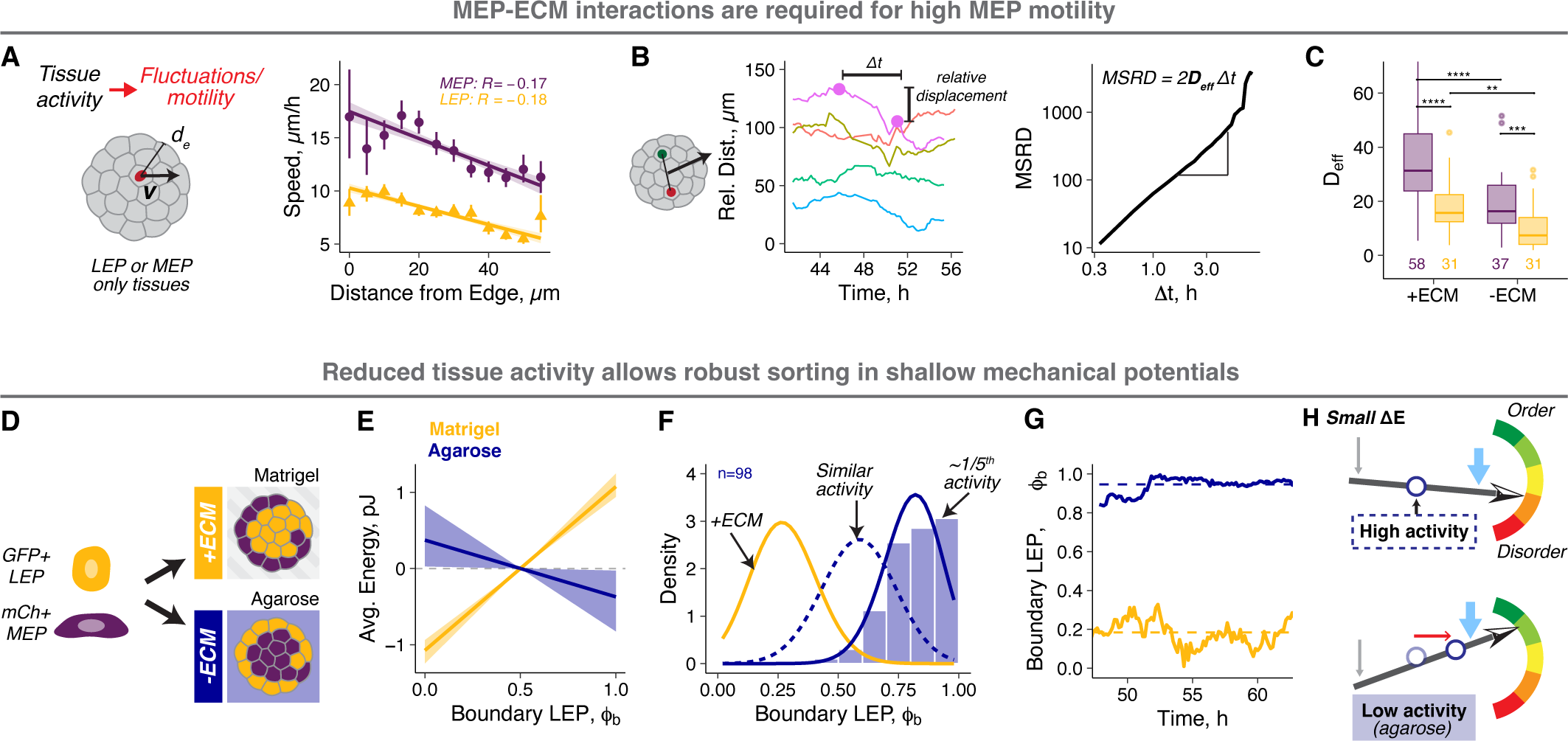
Tissue activity sets the balance between the mechanical potential and macrostate degeneracy. A. Tissue activity is a measure of the kinetic component of the internal energy of tissues and is associated with cell motility. Cell speeds were measured by tracking cell nuclei in LEP- or MEP-only spheroids using time lapse microscopy (n=11 and 14 respectively). Speeds for MEP(purple) and LEP (gold) as a function of distance from the tissue boundary are shown. The Pearson’s correlation coefficients for linear regression are shown. Average speeds and their 95% confidence intervals are represented by the points and error bars respectively. B. The effective diffusion coefficients for cells in spheroids were calculated from the trends for the relative distance between cell pairs. This approach eliminates confounding dynamics from whole organoid movements. The left graph shows example traces of relative distance between cell pairs over time for a representative MEP spheroid. The change in relative distance (relative displacement) was calculated for different time intervals (Δt) and averaged across all times and cell pairs to get the mean squared relative displacement (MSRD). The MSRD vs Δt curves were used to estimate the effective cellular diffusion coefficients (D_)ff_) for each organoid (**Supplemental Information**). C. Effective diffusion coefficients for LEP (gold) and MEP (purple) in the presence and absence of ECM interactions (in Matrigel and agarose microwells, respectively). The lines and hinges for boxplot show the median and the 1^st^ and 3^rd^ quartiles. The number of spheroids analyzed is noted at the bottom of the graph. Asterisks represent the significance of difference between conditions, as follows ns: p > 0.05; *: p < 0.05, **: p < 0.005; ***: p < 0.0005 based on Wilcoxon test. D. Equal proportions of GFP+ LEP and mCh+ MEP were aggregated and cultured in Matrigel (high activity) or agarose microwells (low activity). E. The macrostate energy calculations for organoids in Matrigel (gold) and agarose (navy) using the BCC lattice model. F. The histogram shows probability density for organoids cultured in agarose. The gold line is the fit for organoids in Matrigel (+ECM), and the navy dotted line is the theoretical prediction based on ΔE for agarose with no change in activity. The solid navy line is the theoretical fit to the measured distribution, predicting 5-fold lower activity in agarose compared to Matrigel. G. Structural fluctuations in ϕ_b_over time for representative mammary organoids in Matrigel and agarose (gold and navy respectively). Dashed line is the average of ϕ_b_ for the corresponding condition.

### Engineering the structural ensemble by programming the mechanical potential and activity

This model provides a framework for systematically engineering the ensemble structure by addressing cellular processes like motility and adhesion, and microenvironmental features such as tissue size, composition, and geometry. We first focused on engineering the mechanical potential, which emerges from cell type-specific differences in cell-cell and cell-ECM interfacial tensions. Therefore, we can alter ΔE by perturbing interfacial tensions. For example, the depletion of talin1 (TLN1) and p120 catenin (CTNND1) in MEP (**Supp. Fig. 5A**) increased the **γ**_MEP-ECM_ and **γ**_MEP-MEP_, respectively (**Fig. 5A, Supp. Fig. 5B**). The impact of these perturbations on structural distributions is further regulated by tissue activity. Therefore, to test if programming the mechanical potential and activity predictably alter the structure of organoid ensembles, we prepared spheroids containing equal numbers of normal MEP and TLN1- or CTNND1-knockdown (KD) MEP (tagged with mCh and GFP respectively) (**Fig. 5A**) and compared model predictions with their structural distribution and motility in Matrigel and agarose (**Supp. Fig. 5C**). In Matrigel, normal MEP have high motility and we therefore expect high tissue activity. The estimated ΔE for TLN1-KD spheroids were comparable to that for mammary organoids, while CTNND1-KD MEP spheroids had a small but negative ΔE (**Fig. 5B***, top*). Modeling the impact of these mechanical potentials on high activity tissues predicted robust exclusion of TLN1-KD MEP from the spheroid boundary, but little or no sorting for CTNND1-KD MEP (**Fig. 5C***, top*). In agarose, in contrast, we expect normal MEP to have low motility and we expect correspondingly low tissue activity. The estimated ΔE was small and negative for both TLN1-KD MEP and CTNND1-KD MEP spheroids (**Fig. 5B***, bottom*). Modeling the impact of these mechanical potentials in the context of a 5-fold lower activity predicted increased boundary enrichment for CTNND1-KD MEP in agarose compared to Matrigel (**Fig. 5C***, bottom*). We tested these predictions and observed a remarkable agreement between experiment and theory (**Fig. 5D**).

**Figure 5:**
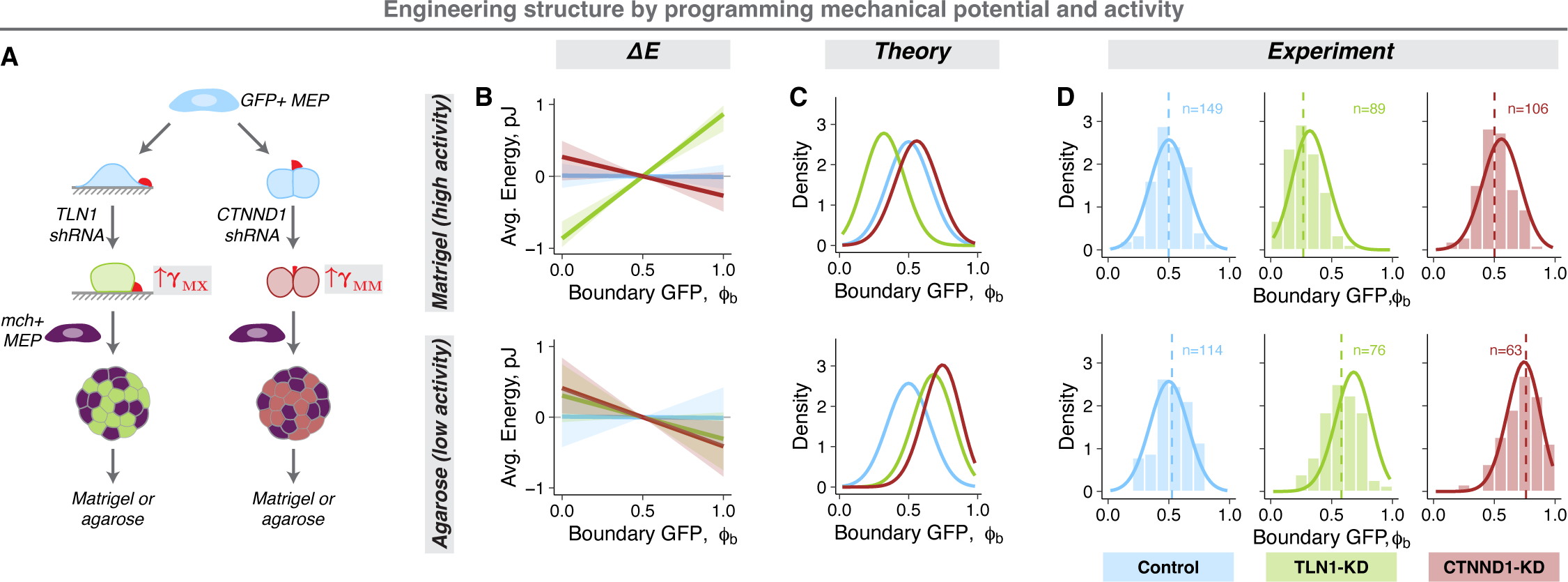
Engineering the structural ensemble by programming the mechanical potential and activity. A. Experimental workflow: The MEP-ECM or MEP-MEP interfacial tensions were perturbed using shRNA against TLN1 (green) and CTNND1 (red). A non-targeting shRNA was used as control (blue). Equal proportion of mCh+ normal and GFP+ shRNA-transduced MEP were aggregated into spheroids (KD-MEP spheroids) and cultured either in Matrigel or agarose. B. The macrostate energy calculations for KD-MEP spheroids in Matrigel (top) and agarose (bottom) using the BCC lattice model. C. The predicted ensemble distributions for KD-MEP spheroids cultured in Matrigel (top) and agarose (bottom). D. The measured probability densities for KD-MEP spheroids cultured in Matrigel (top) and agarose (bottom). Histograms show the distribution of experimental data, dashed vertical lines are the average ϕ_b_, and the solid curves are the theoretical predictions for each condition. The number of observations is noted at the top of the graphs.

### Engineering the structural ensemble by programming macrostate degeneracy

The number and distribution of microstates across macrostates, **Ω**, depends on the cell type proportion (LEP fraction: Φ), tissue geometry (surface area to volume ratio) and the total number of cells (N) (**Supplemental Information**) – all of which can be perturbed quantitatively in organoids. For MEP spheroids with increasing proportion of GFP+ MEP (**Fig. 6A**), the average ϕ_b_increased and matched Φ, in agreement with the theory which predicts that **Ω** is maximum for ϕ_b_ = Φ (**Fig. 6B,C**, *top*). For mammary organoids with varying Φ, ΔE (**Supp. Fig. 6A**) and activity (**Supp. Fig. 6B**) should remain constant, but the average ϕ_b_ should increase due to the underlying shift in **Ω** (**Fig. 6B***, bottom*). Remarkably, the model again accurately predicted the shift in the mean structure across these conditions (**Fig. 6C***, bottom*).

**Figure 6:**
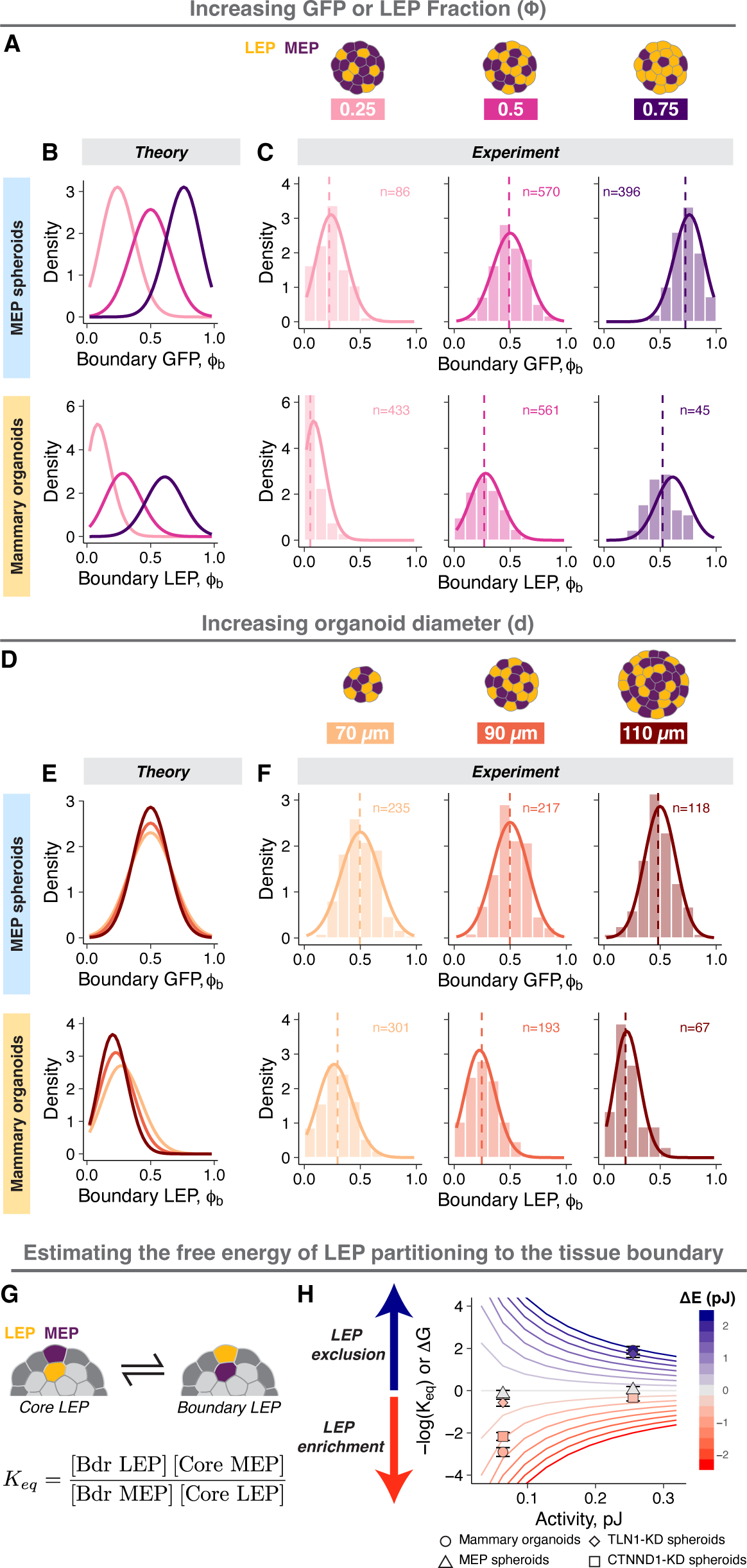
Engineering the structural ensemble by programming macrostate degeneracy. A. Engineering structure by varying LEP proportion: the proportion of GFP+ LEP in mammary organoids or GFP+MEP in MEP spheroids was varied. Tissues with Φ = 0.25, 0.5 or 0.75 (light pink, magenta, and dark purple respectively) were generated. B. Theoretical predictions for MEP spheroids (top row) and mammary organoids (bottom row) with varying Φ. C. The measured probability densities for MEP spheroids (top row) and mammary organoids (bottom row) with varying Φ. Histograms show the distribution of experimental data, dashed vertical lines are the average ϕ_b_, and the solid curves are the theoretical predictions for each condition. The number of observations is noted at the top of the graphs. D. Engineering structure by varying tissue size: the total number of cells per organoid was varied by changing the tissue diameter. Tissues with average diameter of 70 µm, 90 µm, and 110 µm were generated (light orange, orange, and brown respectively). The cell proportions were held constant (Φ = 0.5). E. Theoretical predictions for MEP spheroids (top row) and mammary organoids (bottom row) with varying size. F. The measured probability densities for MEP spheroids (top row) and mammary organoids (bottom row) with varying size. Histograms show the distribution of experimental data, dashed vertical lines are the average ϕ_b_, and the solid curves are the theoretical predictions for each condition. The number of observations is noted at the top of the graphs. G. The equilibrium constant (K_eq_) for the partitioning of LEP between the tissue core to the boundary was calculated from the average occupancy of LEP and MEP in the tissue boundary and core. The free energy change (ΔG) associated with cell translocation is proportional to - log(K_eq_) and determines the favorability of cell translocation. H. Calculations of ΔG for different mechanical potentials and activities in tissues with a diameter = 80 µm containing equal number of LEP and MEP. The contour lines are predictions from the model and are colored by the value of ΔE. Estimated values of ΔG for different experimental conditions are also shown, where points and error bars are the average and standard deviations. The symbols represent different conditions (○: mammary organoids, Δ: MEP spheroids, ◇: TLN1-KD spheroids, □: CTNND1-KD spheroids), and the points are colored by their calculated ΔE.

We also perturbed **Ω** by changing the organoid size (**Fig. 6D**), where an increase in the number of possible sites within the tissue increases the degeneracy of macrostates close to ϕ_b_ = Φ. This had the effect of tightening probability distributions due to the larger difference in **Ω** between adjacent macrostates (**Fig. 6E**). Indeed, both larger MEP spheroids and mammary organoids had narrower distributions (**Fig. 6F**). For the mammary organoids, the larger tissue size also increased ΔE (**Supp. Fig. 6C**), as more LEP must be added to the boundary for the same fractional increase in ϕ_b_. Consequently, larger mammary organoids were additionally shifted toward a lower average ϕ_b_ (**Fig. 6F***, bottom*). Larger tissue size did not change the motility of cells at the tissue boundary, implying similar activity (**Supp. Fig. 6D**). Taken together, these experiments identify the critical parameters in the tissue microenvironment (e.g., cell proportions, tissue size, and ECM) and in single cells (e.g., interfacial tensions and cell motility) that contribute to the average and variance of the structural ensemble. Therefore, quantitative manipulation of these parameters provides a means of systematically engineering the structural ensemble.

### The relative scale of mechanical potential and activity determine the extent of structural order

Analogous to thermodynamic systems, the partitioning of LEP between the core and the boundary is determined by the free energy change (ΔG) associated with their translocation between these two positions. In the absence of a mechanical potential (e.g., MEP spheroids), ΔG = 0 and there should be no LEP enrichment in either compartment (maximum disorder) (**Fig. 1E**). In contrast, a large absolute ΔG (e.g., mammary organoids in Matrigel or agarose) should lead to strong LEP enrichment in either the core or the boundary (high order) (**Fig. 1E**, **4F**). Like thermodynamic state variables, the ΔG for tissues can be inferred from quantitative relationships between tissue state variables such as average total surface energy, average cell dynamics and cell proportions (analogous to enthalpy, temperature, and concentration, respectively). To explore this idea, we calculated an effective equilibrium constant (K_eq_) for LEP translocation from the core to the boundary (**Fig. 6G**), where -log(K_eq_) is proportional to ΔG, and hence the favorability of enrichment of LEP at the tissue boundary. The predictions of the model once again proved surprisingly accurate. For example, the ratio of ΔE and activity determines the magnitude of ΔG, and therefore, the efficiency of cell sorting (**Fig. 6H, Supp. Fig. 6E**). In contrast, changes to macrostate degeneracy have little effect on ΔG, even though these perturbations had a significant impact on the average and variance of tissue structure (**Supp. Fig. 6F,G**). Extensions of this analysis could help identify unexpected empirical relationships between state variables (analogous to equations of state in thermodynamics), predict structural transitions (analogous to transition state theory), and could be transformative in how we understand the emergence of a variety of other macroscopic properties of tissues from the properties and interactions of their individual cellular building blocks.

## DISCUSSION

An ordered spatial arrangement of cells is universally essential to the function of tissues. It is therefore remarkable that at length scales approaching that of single cells, structure can be quite heterogeneous. This is a poorly understood feature of many tissues, both in vivo and in vitro, that has been attributed to both tissue-extrinsic and tissue-intrinsic factors. Here we reveal that this heterogeneity can emerge intrinsically as a consequence of cell mechanics and dynamics by deriving a quantitative relationship between the driving forces of order and disorder that act during the self-organization of human mammary organoids. We show that mammary organoids behave as a dynamic structural ensemble at the steady state, having a reproducible average and variance. This feature of the system allowed us to apply the principle of maximum entropy to link the probability of observing different structural configurations with three parameters: the tissue’s mechanical potential, the degeneracy of possible cellular configurations in space, and the mechanical energy associated with fluctuations in cell position. We then map these tissue-level state variables to the average properties of single cells and their microenvironment, providing a means to make surprisingly accurate predictions for how perturbations to cells and their interactions alter the distribution of structures at the tissue-scale. We conclude that heterogeneity can emerge spontaneously due to mechanical fluctuations, even in the absence of any extrinsic microenvironmental variability or intercellular heterogeneity in gene expression.

Our analysis offers several important and broadly applicable conceptual insights into the self-organization of tissues. First, it explains why the average structure of many tissues differs significantly from the predictions of existing energy-based models of cell sorting^30,38^. Specifically, this discrepancy is a consequence of entropy, which favors the occupancy of otherwise higher energy tissue configurations because they are more numerous compared to lower energy ordered configurations. This serves to shift the average structure away from what would be expected from the mechanical potential alone. Second, it suggests that in all self-renewing tissues, a certain baseline level of structural heterogeneity should be expected because tissues must engage in homeostatic processes that are known to increase their activity, such as cell motility, cell division, apoptosis, immune cell trafficking, and the endocytosis of material from their environment^59–61,66^.

Statistical mechanics provides a quantitative link between the average properties of non-living systems and the underlying properties of their interacting molecular building blocks. Therefore, an important contribution of this study is that it similarly draws an explicit link between the average macroscopic structure of living tissues and the measurable properties of their interacting cellular building blocks. In doing so, it bypasses the need to measure the microscopic state of every gene in every cell in order to understand the properties of tissues as a whole. While many cell-based models reduce the complexity of ∼20,000 gene products to several dozen parameters reflecting cellular state, interactions, dynamics, and tissue material properties^67,68^, these parameters can be hard to measure or can be difficult to map to concrete cell biology of physical processes. Further, many models build in stochasticity and heterogeneity as random noise of unclear origin^27,69^, whereas the statistical mechanical framework applied here allowed us to map tissue activity back to the motility of cells as they engage with the ECM. Consistently, recent studies have highlighted the importance of cell motility (a driver of activity) in regulating solid-to-fluid transitions in tissues during development and cancer progression^66,69^. These studies also invoke thermodynamic concepts like effective energy, temperature, and pressure^63,70^ to predict tissue-level phase transitions, but leave the quantitative relationship between these variables and their biophysical determinants unspecified. By deriving these quantitative relationships explicitly, as we do here, it is intriguing to speculate whether this approach can be extended to other state variables and their derivatives (analogous to Maxwell’s relations) in order to provide insight into how single cells change their state to tune the location of critical points in tissue-level properties like viscosity, pressure, density, volume, and cell positioning that are known to play essential roles in all aspects of tissue development, homeostasis, and breakdown during disease^66,71,72^.

Our analysis of cell-intrinsic drivers of order and disorder in mammary organoids was facilitated by the stability of cellular phenotypes and the tissue microenvironment over experimental timescales. However, unlike this carefully controlled in vitro system, tissue state variables can vary either spatially or temporally (e.g., changes in composition and gene expression with hormone fluctuations^33^ or in breast cancers^16^) under the influence of signals extrinsic to the tissue. This provides one mechanism for tissues to tune their state variables in order to sharpen or re-shape their distribution of structures during development and disease. For example, while activity prevents tissues from getting trapped in metastable configurations during the early stages of self-organization^26^, it also prevents tissues from converging on the most ordered configurations in the latter stages of self-organization. Thus, a gradual decrease in cellular activity (or conversely an increase in the steepness of the mechanical potential) can serve to “anneal” structure towards more ordered distributions. Consistent with this notion, cell motility is progressively reduced in ducts compared to the terminal endbuds during branching morphogenesis of the murine mammary gland^73,74^ and salivary gland^56^. Analogously, cell proliferation and death can modify tissue composition and size, which contribute to microstate degeneracy^75^. In all these cases, the trajectory of self-organization will evolve dynamically along with the changing state variables, channeling tissues towards new or more ordered configurations.

Cells can also regulate their own identity and properties in response to the intrinsic exchange of signals that occurs as they sample different positions (or niches) within the tissue. This property of cells as living materials is wildly different than classical physical systems that are incapable of information exchange and processing. This would suggest that cell state plasticity and tissue structural dynamics are intimately linked, and the coupling between them can drive tissue systems to evolve in unusual ways, for example during development and the progression of disease. In breast cancer, for example, transformed luminal cells must contact the basement membrane in order to break it down and invade^76,77^, and yet their physical properties are not compatible with a sustained position next to the basement membrane in the tissue. Intriguingly, however, transformed luminal cells that localize to the leading edges of invasive tumors and organoids acquire basal characteristics over several days^77,78^. Thus, contact with the basement membrane or the surrounding stroma may provide a new microenvironment for these cells, potentially reprogramming their biophysical properties in a positive feedback loop that facilitates their invasion. Extending these concepts to development, it is intriguing to imagine the guy wires in the Waddington’s famous landscape of cell states^79^ as emerging directly from, and changing with, the evolving structure of the tissue, consistent with the intricate co-evolution of cell state and tissue structure observed during embryogenesis. Therefore, elaborating the statistical mechanical model proposed here to incorporate the link between cell state and cell position^80^ could provide a more quantitative and physical basis for this hypothetical model that has implications for development, stem cell homeostasis, and progression toward diseases like cancer.

The ability to model tissues as dynamic ensembles using concepts of equilibrium statistical mechanics is at its surface quite surprising, as cells and tissues exist far from equilibrium. However, it is now understood that the mathematical foundations of statistical mechanics are deeply linked to information theory, indicating they are far more generalizable than originally assumed^49^. Thus, so long as a system obeys the principle of “balance,” and is therefore at a steady state, one may assume that its most probable distribution of microscopic configurations is one that maximizes its entropy of states subject to the constraint of macroscopic averages. Accordingly, the tools of information theory and statistical physics have now been successfully applied to several important questions in biology, including population dynamics of organisms^51^, stochasticity in cell fate decisions^81–83^, and the heterogeneity in single cell gene expression^84^ and morphology^85,86^, all of which are far from equilibrium processes. While we apply this concept specifically to the mammary gland here, we reason that the application of these principles to other tissue types is warranted and will reveal new strategies for understanding the outcome of development and disease progression. For example, many broad mechanisms of self-organization, such as cell-sorting, are well conserved across tissue types and species (e.g., prostate^31^, salivary gland^56^, neural tube^55^, cochleal epithelium^87^, and developing embryos^10^), even if the detailed structure and the hierarchy of interfacial tensions that guide them differ^88^. While more quantitative measurements of cell mechanics, activities, and tissue geometries are required to define the appropriate state variables in these other tissues, the framework described here should be equally applicable given a suitable structural metric that scales with tissue surface energy. In principle, this statistical mechanical framework could also be adapted to model structural transitions during other morphogenetic processes such as epithelial folding and branching given that tissue activity allows the system to sample across the relevant structural metric during the folding transition and that the corresponding energetic and geometric constraints can be defined quantitatively^5,29^. More broadly, by directly addressing and engineering heterogeneity as a fundamental property of tissues, similar approaches can greatly improve the reproducibility of organoids^21^ and other lab-grown tissues whose variability remains a major roadblock to their more widespread use^19,89,90^.

## MATERIALS AND METHODS

Detailed descriptions for all experimental, analytical, and computational approaches are provided in the Supplemental Information.

## DATA AND CODE AVAILABILITY

All source data will be made available upon publication. Custom MATLAB and R scripts are publicly available through GitHub as:

Code for image analysis - https://github.com/Gartner-Lab/Organoid_Image_Processing.

Code for lattice models - https://github.com/Gartner-Lab/BCC_Lattice_Model.

## ACKNOWLEDGEMENTS

We thank the members of the Gartner Lab and CZ Biohub Theory Group for their valuable suggestions. We particularly thank Dr. Rob Phillips (Caltech) for guidance on analytical approaches during the early stages of this project. This work was funded by the NCI PS-ON grant (U01CA244109) and Department of Defense Era of Hope Expansion Award (W81XWH-13-1-0221) to Z.J.G., the UCSF Center for Cellular Construction (DBI-1548297), and the Barbara and Gerson Bakar Foundation. Z.J.G. is a Chan Zuckerberg BioHub Investigator. V.S. was supported by the Zena Werb Memorial Fellowship through the Helen Diller Family Cancer Center at UCSF. J.L.H. was supported by the NSF Graduate Research Fellowship. FACS was performed on BD Aria 3, supported in part by HDFCCC Laboratory for Cell Analysis Shared Resource Facility through a grant from NIH (P30CA082103). Most lentivirus particles were generated by the ViraCore at UCSF (https://viracore.ucsf.edu).

## AUTHOR CONTRIBUTIONS

V.S., J.L.H and Z.J.G. conceptualized and designed the experiments and the model. V.S. and J.L.H conducted experiments on reconstituted human mammary organoids, analyzed all the data and developed the computational lattice model. J.L.H wrote the script for the quantification of organoid images. V.S., J.L.H and B.V. developed the analytical model. S.F.S. collected and conducted immunostaining on tissue sections of the human breast. J.C.G. and M.A.L provided patient-derived human mammary epithelial cells and assisted in their culture. D.Y. and G.H. provided feedback on the theory and analytical models. V.S. and Z.J.G. wrote the manuscript. All other authors reviewed the manuscript and provided feedback.

---

**Supplemental Figure 1:**
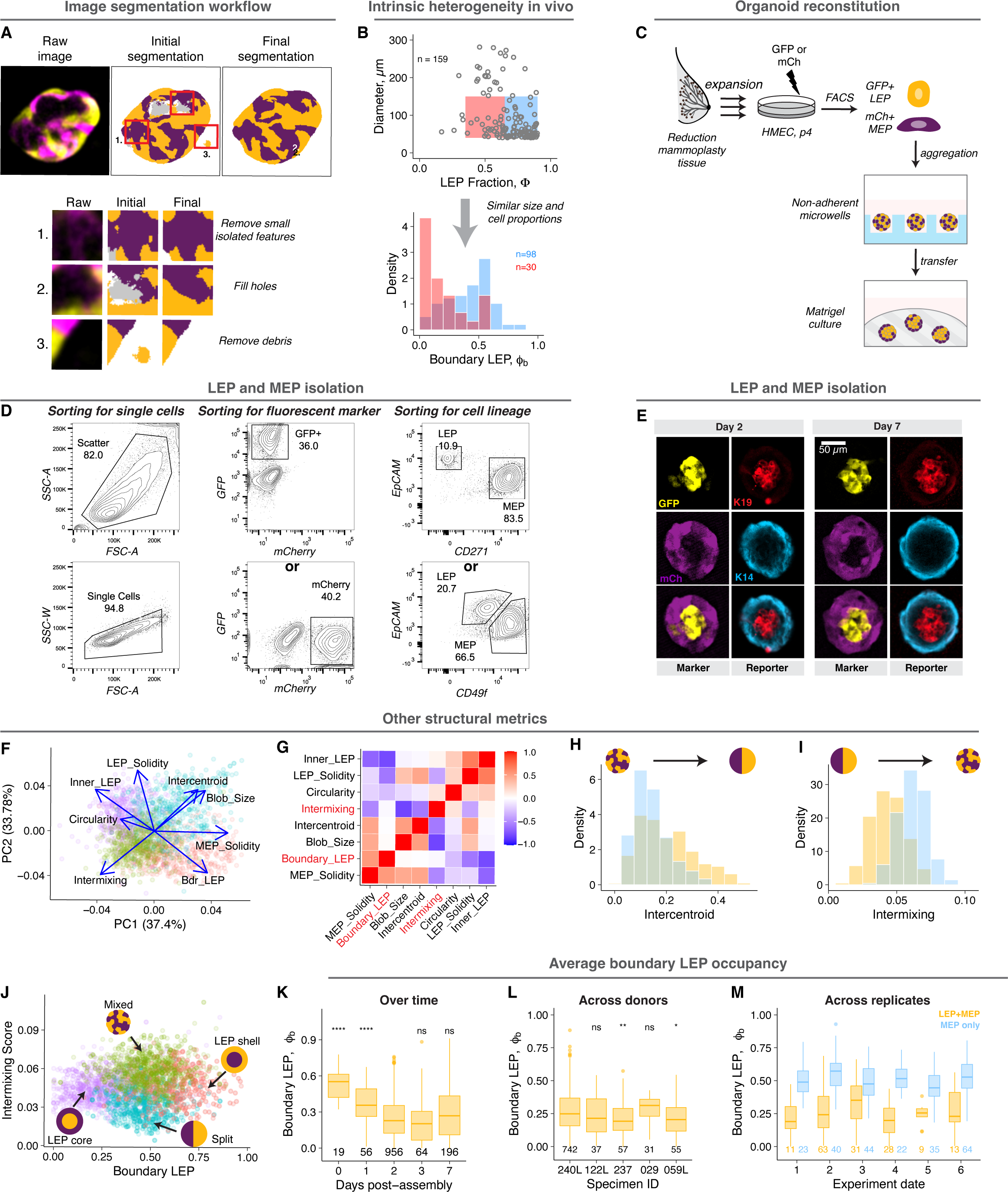
Cell positioning is intrinsically heterogeneous in vivo and in vitro (related to Figure 1). A. Pixel classification and post-processing workflow for image segmentation. Following initial segmentation, images were processed to remove small features, holes, and debris. B. Structural distribution for human mammary gland sections with similar size composition reveals intrinsic heterogeneity in cell position. Data from Fig. 1A was divided by tissue size and LEP proportions. Distributions are shown for tissues smaller than 150 µm in diameter, with LEP proportion between 0.35 and 0.65 (red) or between 0.65 and 0.9 (blue). The number of tissues analyzed in each category are noted on the graph. C. Schematics illustrating the experimental workflow for mammary organoid reconstitution. Fourth passage HMEC were infected with lentivirus expressing cytoplasmic GFP or mCherry and purified by FACS between after 5-7 days. Equal numbers of GFP LEP and mCh MEP were aggregated in non-adherent microwells for 4-6 h prior to transfer to Matrigel. Organoids were imaged two days after reconstitution. D. Cell isolation by FACS. Single cells were isolated based on forward and side scatter intensity. From singlets, GFP+ cells were isolated as GFP+ and mCherry-, while mCherry+ cells were isolated as GFP- and mCherry+. GFP+ LEP were isolated as GFP+, EpCAM-hi and CD271- or GFP+, EpCAM-hi and CD49f-low. mCherry+ MEP were isolated as mCherry+, EpCAM-low and CD271+ or mCherry+, EpCAM-low and CD49f-hi. E. Immunostaining of mammary organoids two and six days after culture in Matrigel. GFP (yellow) and mCh (purple) expression was used as markers for sorted LEP and MEP respectively. The expression of keratin-19 (red) and keratin-14 (cyan) marks luminal and myoepithelial lineages. F. Principal component analysis plot on different structural metrics for organoids containing roughly equal number of LEP and MEP. Organoids were manually annotated as LEP shell, MEP shell, mixed or split (schematics shown in panel g). 500-1000 images were sampled from each category for the analysis. G. Pearson’s correlation matrix for the structural metrics used in the analysis. Boundary LEP fraction and intermixing score were uncorrelated measures of tissue structure. H. The probability density for intermixing score for mammary organoids and MEP spheroids two days post-reconstitution. I. The probability density for the normalized inter-centroid distance for mammary organoids and MEP spheroids two days post-reconstitution. J. Boundary LEP occupancy and intermixing score are effective in separating the 4 annotated structural categories. K. The distribution of LEP boundary occupancy of mammary organoids in Matrigel at different times post-reconstitution. L. LEP boundary occupancy for mammary organoids from different patient specimens in Matrigel, 2 days post-assembly. M. LEP/GFP boundary occupancy for mammary organoids and MEP spheroids in Matrigel across experimental replicates two days post-assembly. Data was collected across multiple experiments. The number of observations is noted at the bottom or top of the graphs. The lines and hinges for boxplots show the median and the 1^st^ and 3^rd^ quartiles. Asterisks represent significance of difference from the reference group (day 2 for panel K, 240L for panel L, and base-mean for panel M), as follows ns: p > 0.05; *: p < 0.05, **: p < 0.005; ***: p < 0.0005 based on Wilcoxon test.

**Supplemental Figure 2:**
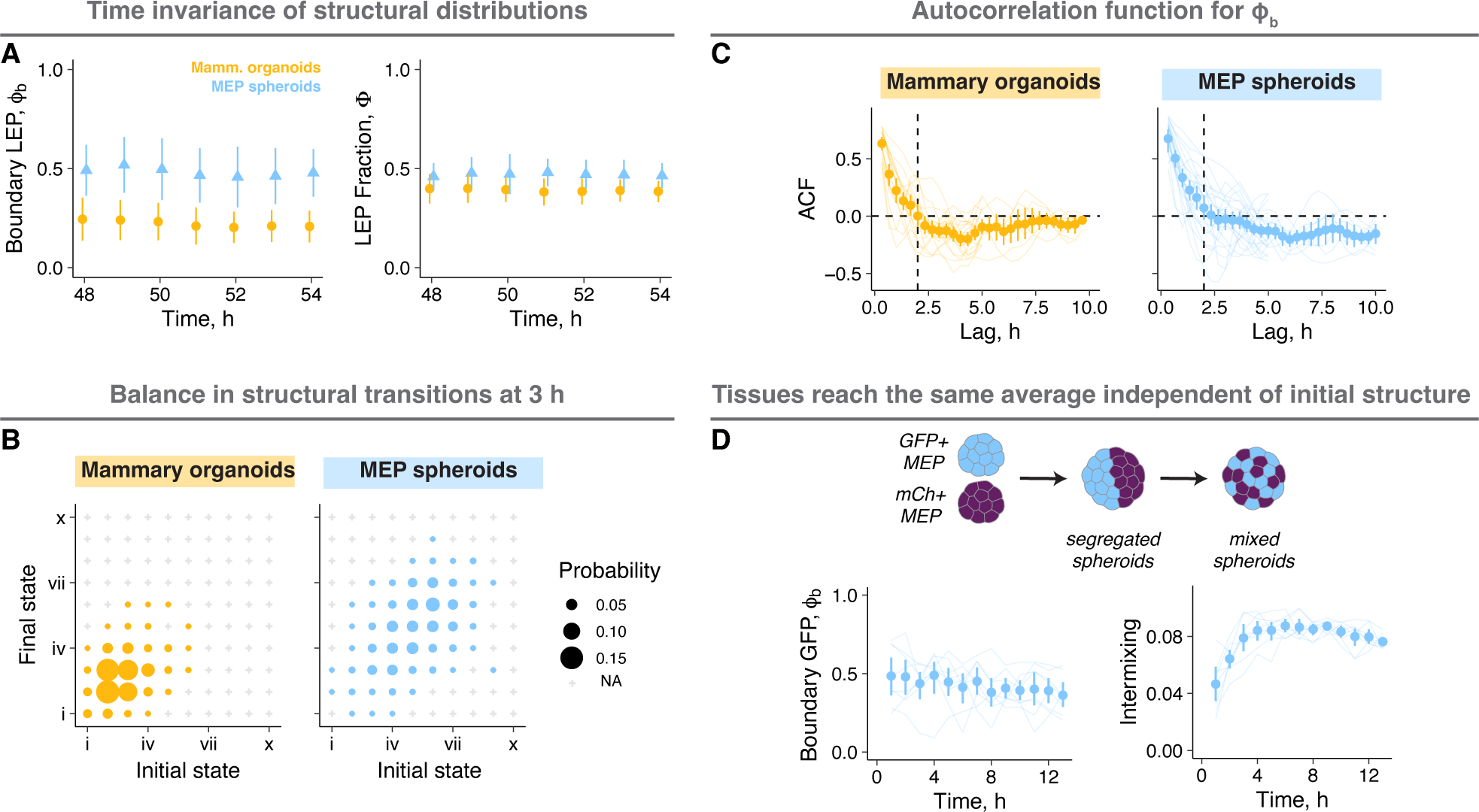
Tissues dynamically sample from the ensemble steady state distribution (related to Figure 2). A. The trends for average ϕ_b_and Φ over time for mammary organoids (gold) and MEP spheroids (blue) analyzed by time lapse microscopy. The whisker plots show the mean and standard deviation for the data. B. The probability of transitioning between any two structural states over a 3-hour window is represented by the size of the circles, similar to Fig. 2C. Any transitions not observed are marked by ‘+’. C. The autocorrelation function (ACF) of ϕ_b_ for mammary organoids and MEP spheroids. The ACF approaches zero after approximately 2 hours (dashed line). The points and error bars are the mean and 95% confidence intervals from binning across all organoids. The lines represent the average for each organoid. D. Small spheroids of GFP- or mCh-were reaggregated to make spheroids with pre-sorted initial structures instead of random structures. There reaggregated spheroids relaxed to random configurations within a few hours, with their average ϕ_b_ and intermixing scores resembling the steady state ensemble average. The whisker plots show the mean and 95% confidence interval for the data.

**Supplemental Figure 3:**
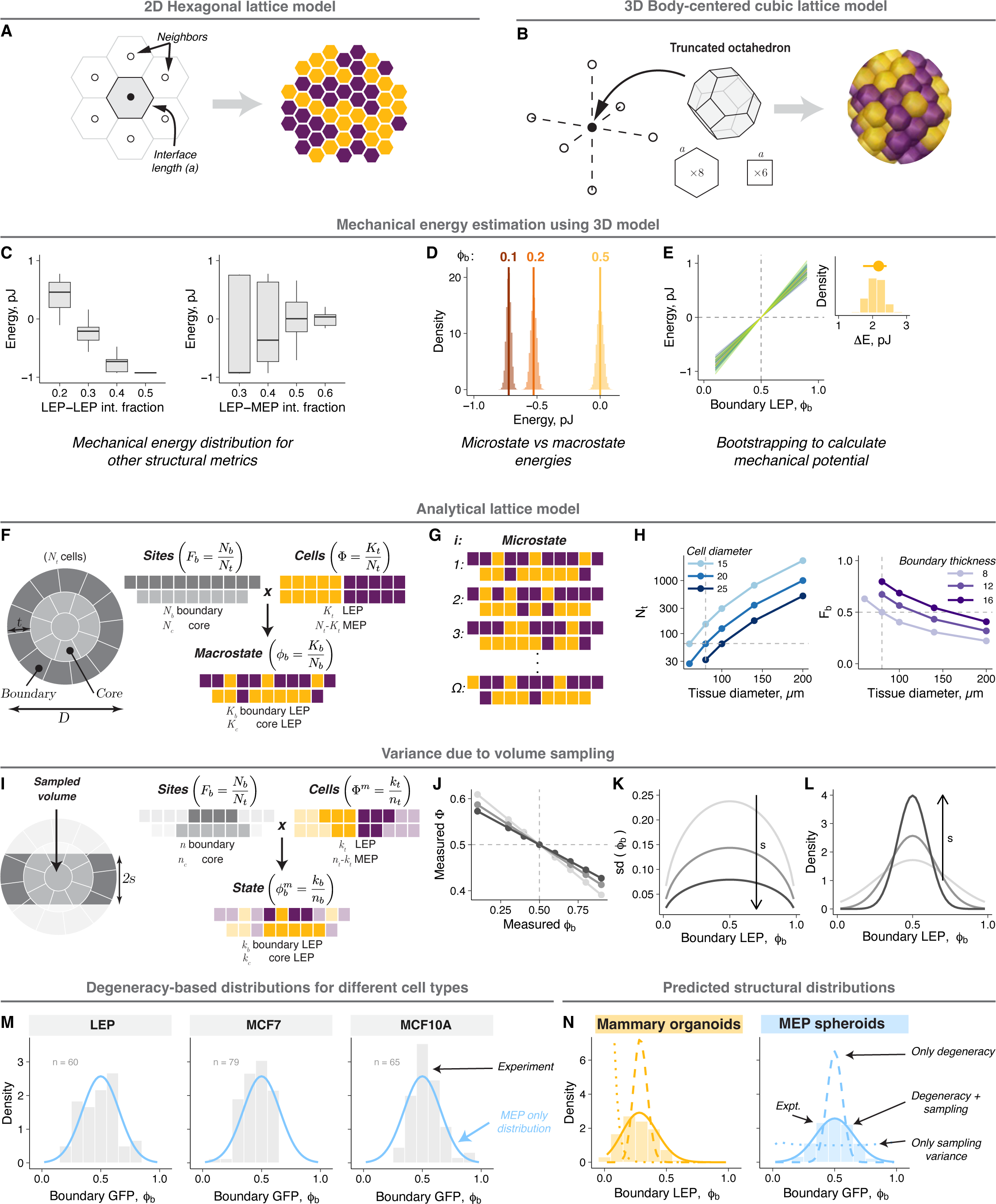
A statistical mechanical framework provides a quantitative description of organoid structural distributions (related to Figure 3). A. A two-dimensional hexagonal lattice model for mammary organoids. Each cell is modeled as a hexagon with 6 neighbors. The cell centers are arranged in a hexagonal grid. B. A three-dimensional body-centered cubic (BCC) lattice model for mammary organoids. Each cell is modeled as a truncated octagon with 6 nearest neighbors (square interfaces) and 8 next nearest neighbors (hexagonal interfaces). The cell centers are arranged in a hexagonal grid. Tissue surface was modeled as a sphere to avoid overestimating the ECM surface area. C. The proportion of cell-cell interfaces (either LEP-LEP or LEP-MEP) do not predict the mechanical energies of the simulated tissues using the BCC lattice. The energies within a macrostate defined by either of these variables are asymmetrically distributed about the average. D. The distribution of microstate energies for the macrostates with ϕ_b_ = 0.1, 0.2 and 0.5. For each macrostate, the energies are symmetrically distributed about the average macrostate energy (vertical line). The difference in energy across macrostates is much larger than that variance within a macrostate. All energies were estimated using the BCC model. E. Bootstrapping was used to obtain confidence intervals for the mechanical potential (ΔE). An example of 50 bootstrapping iterations is shown. The inset shows the distribution of ΔE from 1000 iterations. F. A schematic for the two-compartment lattice model for entropy estimation. Each boundary (dark gray) or core (light gray) lattice site can be occupied by either a LEP (gold) or MEP (purple). Macrostates are defined by the fraction of LEP in boundary (ϕ_b_). G. A schematic representation of structural degeneracy showing many microstates within the same macrostate (e.g., 3 LEP in the boundary). Cells with the same label are identical, and swapping two cells of the same identity yields identical microstates. H. Estimated number of cells and boundary fraction for tissues of different sizes, as a function of the diameter of a single cell and the thickness of the boundary compartment. Based on the average organoid size, tissues with diameter = 80 µm and a boundary thickness of 8 µm were used unless specified otherwise (dashed line). For these parameters, lattice sites were equally distributed between the boundary and the core. I. Schematic showing the effect of sampling the middle tissue section on the measured LEP fraction and LEP boundary occupancy. J. The relationship between the measured LEP boundary occupancy and the measured LEP fraction for different imaging depths in organoids with equal proportion of LEP and MEP. For sorted organoids, the measured LEP fraction is higher than the expected value. K. Estimated standard deviation for ϕ_b_ due to volume sampling for different imaging depths. L. Predicted structural distributions for MEP spheroids resulting from combining structural degeneracy and volume sampling for different imaging depths. M. Experimentally measured structural distributions for spheroids containing equal number of GFP+ and mCh+ LEP, MCF7 or MCF10A cells. The blue line is the fit for the distribution for MEP spheroids. N. Comparison of experimental and predicted structural distributions for mammary organoids and MEP spheroids. Histograms show the experimental distribution. Dashed and dotted lines are predictions based on degeneracy only or volume sampling only. The solid line is the final analytical model prediction.

**Supplemental Figure 4:**
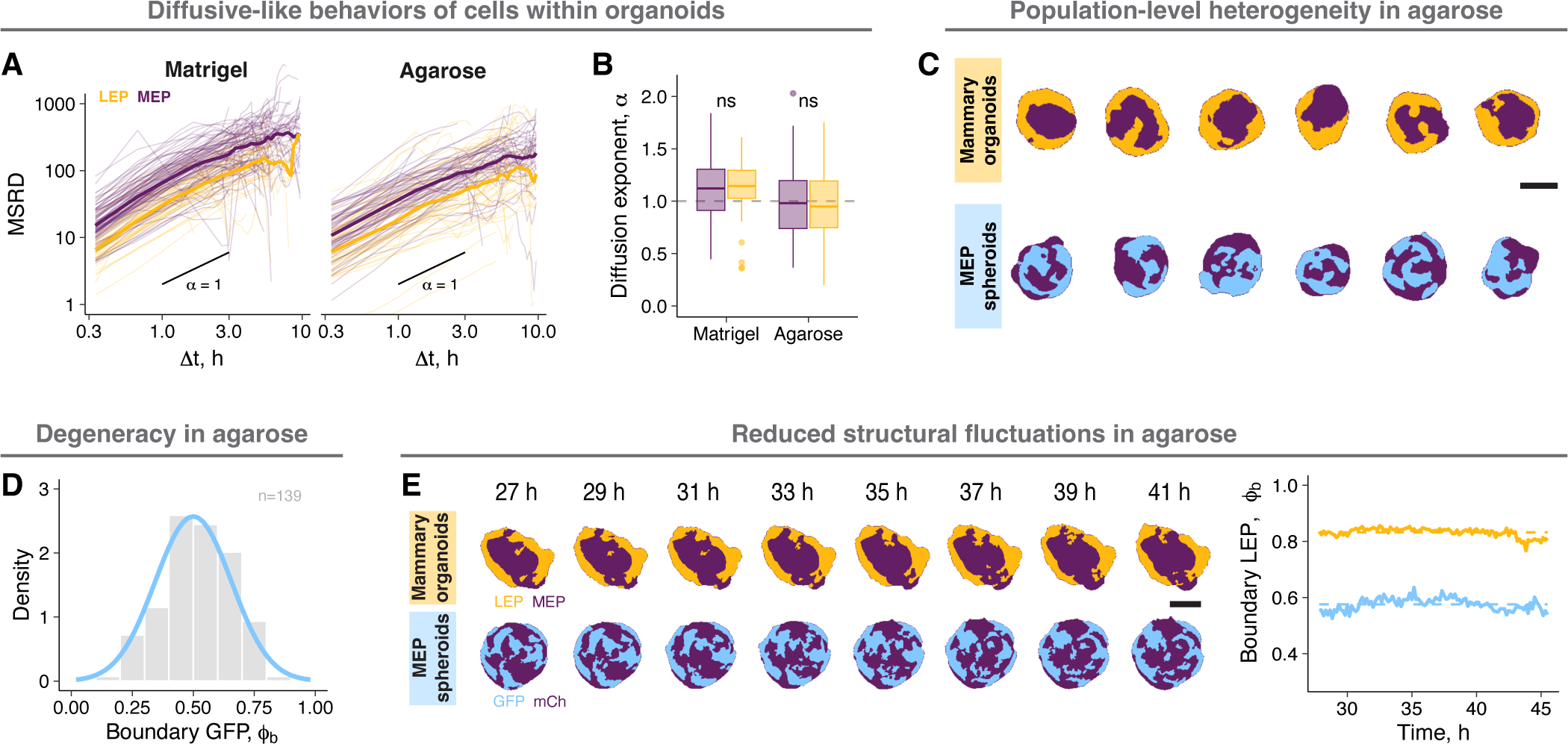
Tissue activity sets the balance between the mechanical potential and macrostate degeneracy (related to Figure 4). A. Example MSRD vs Δt curves for MEP (purple) or LEP (gold) spheroids in Matrigel and agarose. Time intervals less than 3 h we used to fit the data to the diffusion models to calculate D_eff_ and α, as the curves were mostly linear at these time scales. B. The diffusion exponents for MEP (purple) or LEP (gold) spheroids in Matrigel and agarose. The cells show diffusive-like behaviors (α = 1). C. Example segmentations of mammary organoids and MEP spheroids in agarose at steady state (24 h). Scale bar = 50 µm. D. The structural distribution for MEP spheroids in agarose. The blue line is the distribution of MEP spheroids in Matrigel. The number of spheroids is noted on the graph. E. Representative segmented mammary organoids and MEP spheroids cultured in agarose and followed by time lapse microscopy. Scale bar = 50 µm. Graph shows the quantification for the organoids shown.

**Supplemental Figure 5:**
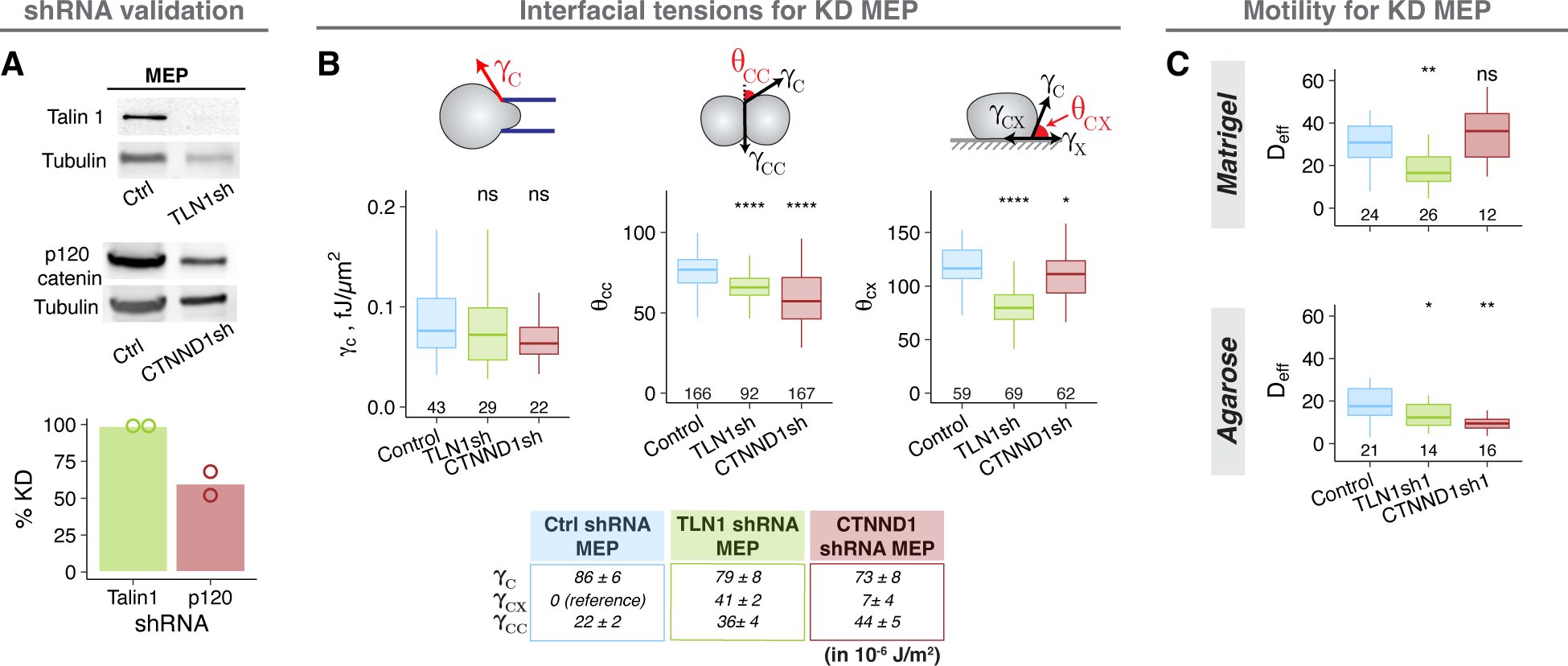
Engineering the structural ensemble by programming the mechanical potential and activity (related to Figure 5). A. Depletion of Talin-1 and p120 catenin protein expression in MEP upon treatment with shRNA against TLN1 and CTNND1 respectively. Top panels show representative western blots for Talin 1 and p120 catenin expression along with α-tubulin expression (loading control). All intensity measurements were normalized to the loading control. The graphs show percent reduction in normalized protein expression in shRNA-treated cells compared to non-targeting shRNA-treated cells. B. Interfacial tension measurements for KD-MEP. Cortical tension, cell-cell contact angles and cell-ECM contact angles are shown using boxplots. Estimated cell-cell and cell-ECM interfacial tensions are listed in the table. C. Apparent diffusion coefficients for KD-MEP in Matrigel (top) and agarose (bottom). The number of spheroids analyzed is noted at the bottom of the graph. Asterisks represent significance of difference from the reference group (ctrl shRNA), as follows ns: p > 0.05; *: p < 0.05, **: p < 0.005; ***: p < 0.0005 based on Wilcoxon test.

**Supplemental Figure 6:**
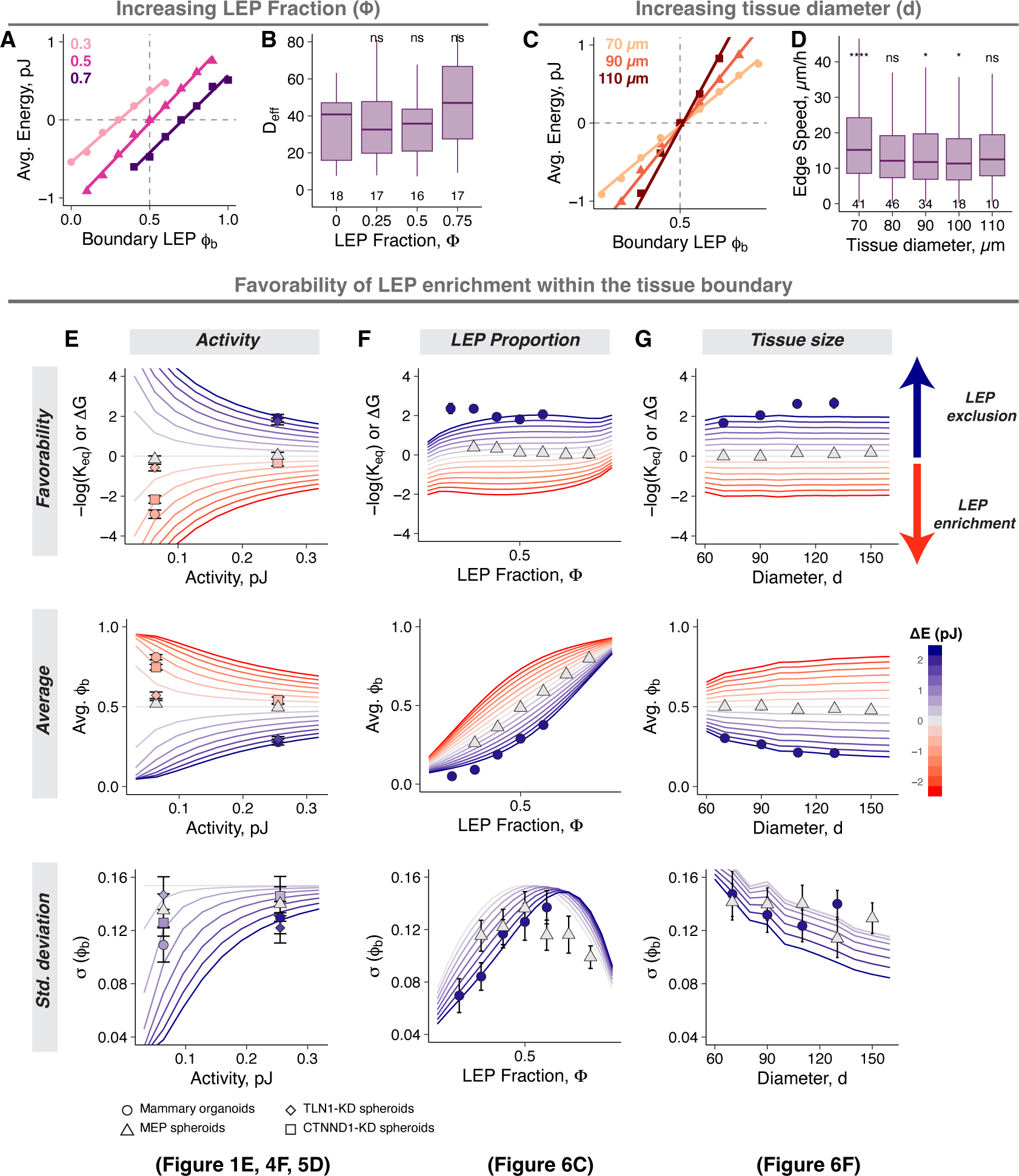
Engineering the structural ensemble by programming macrostate degeneracy (related to Figure 6). A. The macrostate energy calculations for mammary organoids with increasing Φ using the BCC lattice model. B. Apparent diffusion coefficients for MEP in Matrigel for organoids with varying Φ. The number of spheroids analyzed is noted at the bottom of the graph. C. The macrostate energy calculations for mammary organoids with increasing diameter using the BCC lattice model. D. Cell speeds near the tissue boundary for MEP spheroids of varying sizes in Matrigel. The number of spheroids analyzed is noted at the bottom of the graph. E. Calculations of ΔG, average ϕ_b_, and standard deviation of ϕ_b_ for different mechanical potentials and activities in tissues with a diameter = 80 µm containing equal number of LEP and MEP. The lines are predictions from the model and are colored by the value of ΔE. Estimated values for different experimental conditions are also shown, where points and error bars are the average and standard deviations. The symbols represent different conditions (○: mammary organoids, Δ: MEP spheroids, ◊: TLN1-KD spheroids, □: CTNND1-KD spheroids), and the points are colored by their calculated ΔE. F. Calculations of ΔG, average ϕ_b_, and standard deviation of ϕ_b_ for different mechanical potentials and LEP fractions in tissues with a diameter = 80 µm and activity corresponding to Matrigel. Estimated values for different experimental conditions are also shown, where points and error bars are the average and standard deviations. The symbols represent different conditions (○: mammary organoids, Δ: MEP spheroids), and the points are colored by their calculated ΔE. G. Calculations of ΔG, average ϕ_b_, and standard deviation of ϕ_b_ for different mechanical potentials and tissue sizes LEP fraction = 0.5 and activity corresponding to Matrigel. Estimated values for different experimental conditions are also shown, where points and error bars are the average and standard deviations. The symbols represent different conditions (○: mammary organoids, Δ: MEP spheroids), and the points are colored by their calculated ΔE.

