## Supplemental Information for "Configurational entropy is an intrinsic driver of tissue structural heterogeneity"

### Supplementary Information

### 1 Experimental Methods

#### 1.1 Cell culture and isolation

**1.1.1 Human mammary epithelial cells (HMEC):** De-identified normal, finite-lifespan primary HMEC were provided by Drs. Martha Stampfer and James Garbe (Lawrence Berkeley National Laboratory). HMEC were cultured from tissues removed during reduction mammoplasties, and expanded to fourth passage in M87A medium as described previously[1]. HMEC from tissue donated by a 19-year old individual with bilateral breast hypertrophy (240L) were used for most experiments, unless specified otherwise. Key experiments were validated in HMEC from other donors as detailed in **Table 2**. All HMEC were cultured in complete M87A medium with Penicillin-Streptomycin (100 U/mL) at 37°C with $CO_2$ up to 80-90% confluence.

**1.1.2 Additional cell lines:** MCF10A and MCF7 cells were purchased from ATCC. MCF10A cells were cultured in DMEM/F12 + 5% horse serum + 20 ng/ml EGF + 1 μg/ml hydrocortisone + 1 ng/ml cholera toxin + 10 μg/ml insulin. MCF7 cells were cultured in DMEM/F12 + ITS-X. Cells were grown at 37°C with $CO_2$ up to 80-90% confluence.

**1.1.3 Plasmid generation and lentiviral packaging:** pSicoR plasmids containing EF1α promoter-driven GFP or mCherry were obtained from UCSF ViraCore. The H2B-mScarlet construct was made by cloning H2B-mScarlet sequence in the pSicoR vector using the AvrII and EcoRI digestion sites. Validated shRNA constructs for the depletion of TLN1 (TRC Clone ID TRCN0000310369) and CTNND1 (TRC Clone ID TRCN0000333514) in pLKO.1 were purchased from Millipore-Sigma. Alternatively, the same shRNA sequence was cloned into GFP-pSicoR downstream of the U6 promoter using HpaI and NotI digestion sites. All plasmid sequences were confirmed by sequencing.

Concentrated lentivirus was either purchased from UCSF ViraCore or prepared in-house. For lentiviral packaging, HEK293T/17 cells (UCSF Cell Culture Facility) were grown in DMEM + GlutaMAX (ThermoFisher Scientific #10566016) with 10% FBS and 1mM sodium pyruvate, then switched to lentivirus packaging medium (Opti-MEM + GlutaMAX (ThermoFisher Scientific #51985034) with 5% FBS and 1mM sodium pyruvate) 12-16 h before transfections. Cells at 80-90% confluence were transfected with the plasmids for the gene of interest, pCMV-VSV-G and pCMV delta R8.2 using lipofectamine 3000 (ThermoFisher Scientific, #L3000015). pCMV-VSV-G was a gift from Bob Weinberg (Addgene plasmid #8454) and pCMV delta R8.2 was a gift from Didier Trono (Addgene plasmid #12263). Lentivirus-containing medium was harvested at 24 h and 48 h after transfection. The pooled medium was centrifuged at 2000 g for 10 minutes at 4 °C and filtered using 0.45 μm filter (Corning #431220) to remove debris. Lenti-X concentrator (TakaraBio #631231) was added to 1/3 supernatant volume, mixed by inverting and stored overnight at 4 °C. The mixture was centrifuged at 1500 g for 45 minutes and the pellet was resuspended in 1/100th of original volume in M87A medium, and aliquots were stored at -80°C. Titers were calculated by infecting fifth passage MEP and measuring the proportion of GFP+ or mCh+ cells by fluorescence activated cell sorting (FACS) 4-5 days after infection.

**1.1.4 Viral transduction of HMEC:** Fourth passage HMEC were transduced with lentivirus upon thawing at a multiplicity of infection of 1-3 (to target 40-60% transduction efficiency) into a half-volume of M87A medium containing 2 μg/ml polybrene (Millipore-Sigma #TR-1003). After 3 hours, M87A medium was added to full volume. After 24-48h, the virus containing medium was discarded and replaced with fresh M87A medium. For pLKO.1-based constructs, 1 μg/l puromycin was also added at 48h for selection of transduced cells. Cells were grown up to 80-90% confluence (5-7 days). Transduced cells were isolated by FACS based on GFP or mCherry expression. Western blots were used to confirm the expression or depletion of constructs.

**1.1.5 Epithelial cell isolation by FACS:** Isolation of single LEP and MEP was described previously. Briefly, HMEC were detached using 0.05% trypsin, passed through a 40 μm nylon filter, and washed once in M87A medium. The cells were resuspended at $< 5\text{x}10^7$ cells/ml in M87A medium containing Pacific Blue-EpCAM and APC/Cy7-CD49f or APC/Cy7-CD271. Cells were incubated on ice for 20 minutes, washed and resuspended in PBS + 2% w/v bovine serum albumin at $< 10^7$ cells/ml. Cells were sorted on a BD FACSAria III running FacsDiva software using the 4-way purity setting. GFP+ cells were gated as GFP+/mCherry- and mCherry+ cells were gated as GFP-/mCherry+ (**Supp. Fig. 1D**). LEP were gated as $\text{EpCAM}^{high}$ and $\text{CD49f}^{low}$ or CD271- and MEP were gated as $\text{EpCAM}^{low}$ and $\text{CD49f}^{high}$ or CD271+ (**Supp. Fig. 1D**). Cell purity was assessed with immunofluorescence by staining for keratin 19 and keratin 14 as luminal and myoepithelial markers respectively. A complete list of antibodies used is provided in **Table 2**. For specimens with LEP proportion <5%, LEP enrichment was performed using the Easy- Sep™ Human CD271 Positive Selection Kit II (StemCell Technologies #17849). Cells were incubated with CD271-selection cocktail, FcR blocker, Pacific Blue- EpCAM and APC/Cy7-CD49f for 15 min prior to the addition of magnetic nanoparticles for 15 min. The cell suspension was incubated on the EasySep magnet for 10 min before transferring the supernatant to a new tube. The transferred cells were resuspended in FACS buffer and sorted as described above. The details for all antibodies used provided in **Table 3**.

#### 1.2 Organoid reconstitution and imaging

**1.2.1 Photolithography and PDMS stamps for agarose microwells:** Custom photomasks were ordered from CAD/Art Services, Inc to make arrays of 120 or 180 μm diameter microwells for organoid reconstitution and 20x40 μm oblong microwells for cell-cell contact angle analysis. Silicon masters were prepared using photolithography as described[3]. SU-8 UV-curing resin (MicroChem) was spin-coated onto pre-cleaned silicon wafers using a spin coater. SU-8 2050 was spun for 30 seconds at 1000 rpm to achieve 200 μm-high features, while SU-8 2025 was spun at 4000 rpm for 20 μm-high features. The wafer was baked at 95 °C to remove excess resist solvent, then exposed under a

365-nm UV lamp with a printed photomask for approximately 6 min for thick features and 2 min for thin features. After baking at 95 °C for 1 min, cooled wafers were submerged in PGMEA developer at 2 cm depth on a shaker until the uncured resist was dissolved away. Excess resist was rinsed off with PGMEA developer, then the wafer was rinsed with isopropanol, dried, and baked for a few minutes at 95 °C to remove excess solvent. SYLGARD™ 184 Silicone Elastomer (Dow Corning #4019862) polymer and crosslinker at a 10:1 ratio were mixed and poured onto the silicon master, degassed in a vacuum desiccator, and baked at 60°C overnight. The PDMS stamps with the array patterns were cut into 0.5 or 1 cm squares to fit 24-well plates and stored in 70% ethanol. Agarose microwells were cast from 3% w/v high-melting point agarose (AllStar Scientific 490-050) in PBS using the patterned PDMS stamps.

**1.2.2 Mammary organoid reconstitution:** Sorted HMEC were reconstituted into organoids by aggregating cells in suspension[2] (**Supp. Fig. 1C**). Unless specified otherwise, sorted GFP+ LEP and mCherry+ MEP were combined in M87A media at a 1:1 ratio at a total cell concentration of 300,000 cells/ml. 150 μL of the cell suspension was added onto agarose microwells in a 24-well plate and centrifuged twice at 160 g in 4 °C with moderate ramp acceleration and deceleration in a swinging bucket rotor centrifuge, switching the plate direction after the first spin to distribute cells evenly across the microwells. Excess cells were removed by washing with M87A medium. Alternatively, cells were centrifuged into 96-well cell repellant plates (Greiner Bio-one #650970) or EZsphere plates (Nacalai USA #4860-900SP). The cells were incubated at 37°C till they formed cohesive aggregates, typically 4-6 hours. For self-organization in ECM hydrogels, 24-well glass bottom plates (Greiner Bio-one #662892) were coated with 10-20 μL cold growth factor-reduced Matrigel (Corning #354230) and placed at 37 °C for 20 minutes. The aggregates were gently dislodged by pipetting using a wide bore pipette, and collected by centrifugation at 10 g for 30 s with low acceleration and deceleration. The aggregates were resuspended in 40 μL Matrigel and added to the Matrigel-coated wells. Matrigel was allowed to solidify for 30 minutes at 37 °C before adding medium. For self-organization in agarose, the cell aggregates were cultured in the agarose microwells prior to imaging.

**1.2.3 Confocal microscopy of reconstituted organoids:** Organoids were imaged on either a spinning disk confocal microscope (Zeiss Cell Observer Z1 equipped with a Yokagawa spinning disk) or a laser scanning confocal microscope (Zeiss LSM800) running ZEN software using a Zeiss LD 20x/NA 0.4 air or LD 40x/NA1.1 water objective. Seven to 11 z-stacks spaced 5 μm apart were acquired for each organoid. All imaging was performed under environmental control with 5% $CO_2$ at 37 °C. For steady state analysis, organoids in Matrigel or agarose were imaged two or one day after reconstitution respectively. For time-lapse analysis, images were acquired at 10 or 20 minute intervals for 8-16 hours. Longer time lapse acquisitions were not performed to maintain viability and minimize phototoxicity, as extended imaging resulted in poor organization. All images were collected when tissues were expected to be at steady state (45 h and 24h for organoids in Matrigel and agarose respectively) unless specified otherwise. Organoids near the edge of the Matrigel dome and bubbles, or those touching the coverglass were not included in the dataset.

**1.2.4 Reducing variability of mammary organoids:** We reconstituted organoids from finite passage, lineage-restricted LEP and MEP that have minimal potential to differentiate into other cell types over the course of the experiments (<7 d). These cells maintain distinct identities, yet have relatively low intra-lineage heterogeneity compared to freshly isolated primary cells[2].The numbers and proportions of cells within organoids were controlled by mixing purified single cell suspensions (LEP and MEP) at desired concentrations. As some variability is expected due to Poisson loading of the microwells, we used additional filters to only analyze organoids with similar size, cell proportion and circularity (section **??**). Further, HMEC growth and self-organization is sensitive to handling technique, batch-to-batch variability in the media additives and other environmental factors. Therefore, we used the intra-day average structure of normal mammary organoids as an additional selection criterion, where only experiments with <60%correctly sorted organoids were analyzed.

#### 1.3 Organoid image segmentation and structural analysis

**1.3.1 Manual annotation:** Organoid z-stacks were qualitatively categorized as LEP core, mixed, split or LEP shell, using a random channel-swapping approach to blind the user to the true identity of LEP and MEP pixels. These annotations were used to ensure equal representation of all structural categories in the training data. Organoid classification also helped identify the relevant quantitative metrics to distinguish between the structural categories (**Supp. Fig. 1d-f**). During the annotation process, the user also marked the center z-section (based on cross-sectional area) to use in automated image quantification.

**1.3.2 Image pre-processing:** The three z-slices at the center of the organoid were isolated from each z-stack and prepared for segmentation by background subtraction and automatic contrast adjustment in FIJI. A Gaussian blur was applied to images acquired on the LSM to reduce the pixel noise. Channels were standardized by swapping where reasonable, so that LEP were always in channel 1. This helped remove any bias due to expression or imaging artifacts as the fluorescent marker should not affect the self-organization process.

**1.3.3 Image segmentation:** Organoid cross-sections were segmented using the pixel classification module in Ilastik[4], based on a Random Forest classifier. The training datasets were generated from random samples of images from different structural categories to ensure their even representation. The training dataset was sparsely annotated into four pixel categories: MEP, LEP, ECM, and hole. The hole category denotes regions within the organoid with very low signal, such as lumens, vacuoles, or the interiors of large organoids where light penetration is weak. Two-dimensional filters of different sizes were applied to background-subtracted two-channel fluorescence data. The training data was annotated until regions of uncertainty were only localized to boundaries between pixel classes. A single pixel classification pipeline was used to batch process all organoid images with similar imaging quality. Separate pixel classification pipelines were generated for each of two microscopes (spinning disk confocal and laser scanning confocal) and two levels of fluorescent protein expression (bright and dim) to account for image variation between microscopes and constructs.

**1.3.4 Image post-processing:** Segmented images were exported and processed using a custom script in MATLAB. Due to natural variations in fluorescence and cell morphology, some regions are very difficult to segment either manually or automatically. Processing was used to smoothen the border of the organoid and reasonably infer unassigned or ambiguous pixel identities. It removed labeled areas outside of the main organoid, removed small speckles, reassigned internal holes as either cells or ECM by interpolation, and used a disk structuring element to smooth out crevices in the organoid that can be replaced with cell pixels (**Supp. Fig. 1A**). The majority of images required minimal processing, unlike the more challenging partially lumenized LEP only spheroids (example shown in **Supp. Fig. 1A**).

**1.3.5 Image quantification** : The morphological parameters of each segmented slice were calculated using a custom script in MATLAB. A complete list of variables, their purpose, and their derivations are provided in **Table ??**. Each pixel was adjacent to its 8 neighbors (Chebyshev distance 1). Where applicable, absolute lengths and areas were converted from pixels to microns or microns squared respectively. The LEP fraction ($\Phi$) was defined as

$$\Phi = \frac{\text{LEP pixels}}{\text{Total pixels}} \,. \tag{1}$$

The LEP boundary occupancy ($\phi_{\text{b}}$) was defined as

$$\phi_{\text{b}} = \frac{\text{LEP pixels in boundary}}{\text{Total pixels in boundary}} \,. \tag{2}$$

For MEP only spheroids or organoids containing GFP+ KD-MEP, the GFP channel was assigned to the LEP channel during pixel classification. In those cases, $\Phi$ and $\phi_{\text{b}}$ measured the GFP fraction and GFP boundary occupancy respectively.

Unless specified otherwise, organoids that met the following criteria were used for analysis:

- organoid circularity > 0.5,
- $\Phi$ was between 0.4 and 0.6,
- the tissue diameter was between 70 and 120 $\mu\text{m}$,
- the average $\phi_{\text{b}}$ of all normal organoids imaged on that day was less than 0.4 (the mean plus standard deviation for the population ensemble).

**1.3.6 Time lapse analysis:** The image segmentation and quantification pipeline was modified slightly to analyze time lapse data. The center slice was assumed to be the slice with the largest area; in the case of a contiguous near-tie (within 15% of the largest area), the center slice among those was selected. Only organoids that met all of the following criteria were analyzed.

- imaging was started after 45 h or 22 h for organoids in Matrigel or agarose respectively, when tissues are expected to be at steady state,
- the average $\Phi$ was between 0.35 and 0.65,

- the standard deviation of $\Phi$ was less than 0.1,
- the tissue diameter was between 70 and 140 $\mu m$,
- the standard deviation in in tissue diameter was less than 10% of the average diameter, and
- the average $\phi_b$ of all organoids imaged on that day was less than 0.4 (the mean plus standard deviation for the population ensemble).

*Structural balance calculations:* Macrostates were assigned to all organoids at each time point based on the instantaneous $\phi_b$ (10 equal bins centered at 0.05, 0.15, ..., 0.95) (**Fig. 2C**). For different time intervals ($\Delta t$) between 20 min to 3 hours, the transition probability between any two macrostates was calculated across all organoids. For the system to obey the laws of balance, there should be no net flux between any two macrostates, i.e. $p_{i \to j} = p_{j \to i}$ for all $i$ and $j$. This can be described as a probability matrix where the rows and columns are the initial and final structural macrostates, and the matrix values are the transition probabilities. A system obeying detailed balance corresponds to a diagonally symmetric matrix.

*Structural relaxation calculations:* The average $\phi_b$ for all organoids across time was calculated ($\overline{\phi_b}$) At each time point, tissues were assigned bins (deviation states: -0.2, -0.1, 0, 0.1, 0.2) based on the value of $\phi_b - \overline{\phi_b}$ (**Fig. 2D**). Any tissues with deviations > 0.25 from the mean were excluded. The measurements were grouped by their deviation states, and the average structure at time intervals between 20 min and 5 hours was calculated and plotted. For the system to be at steady state, while transient deviations away from the population mean occur, tissues on average should relax towards the population average at sufficiently long times.

*Autocorrelation analysis:* The autocorrelation function was calculated for each organoid and different lag times ($\Delta t$) as

$$ACF_{\Delta t} = \frac{\sum_0^{T-\Delta t}(\phi_{b,\,t} - \overline{\phi_b})\,(\phi_{b,\,t+\Delta t} - \overline{\phi_b})}{\sum_0^{T}(\phi_{b,\,t} - \overline{\phi_b})^2}, \tag{3}$$

where $\phi_{b,\,t}$ and $\phi_{b,\,t+\Delta t}$ are the LEP boundary occupancy at times $t$ and $t + \Delta t$ respectively, and $T$ is the maximum time for acquisition.

#### 1.4 Structural quantification for human tissue sections:

Details for sample collection and staining were described previously[5]. Tissue sections were stained for keratin-19 (K19) and keratin-14 (K14), and imaged using a Nikon Eclipse Ti2 microscope. LEP and MEP were classified as K19+ and K14+/K19- respectively. Images from normal tissue specimens were segmented and quantified using a similar approach as that used for the mammary organoids.

#### 1.5 Interfacial tension measurements

**1.5.1 Micropipette aspiration:** Cellular cortical tension was measured as described previously[6]. Sorted cells were resuspended at 5000 cells/ml, kept on ice till ready to use (up to 5 h), and warmed to room temperature prior to aspiration. A cell was aspirated into an 8-11 $\mu m$ diameter glass micropipette at a beginning pressure of 0.03 kPa, and pressure was increased in 0.03 kPa steps till the cell was sufficiently deformed. At each pressure, after waiting for 30 seconds, three images were taken using a 40x air objective on a Zeiss Axiovert 200M running SlideBook software. The average cellular deformation inside the pipette ($L_p$) was measured in FIJI, and critical pressure ($\Delta P_{\text{crit}}$) was identified as the aspiration pressure where $L_p$ equals the pipette radius ($R_p$). Any cells that blebbed were discarded. The cortical tension ($\gamma_{\text{c}}$) was calculated as follows, where $R_c$ is the radius of the cell

$$\gamma_{\text{c}} = \Delta P_{\text{crit}} / \left( \frac{1}{R_p} - \frac{1}{R_c} \right) . \quad (4)$$

**1.5.2 Cell-ECM contact angles:** Cell-ECM contact angle measurements were described previously[2]. Eight-chambered coverglass (Nunc Lab-Tek II, ThermoFisher Scientific #155409) were coated with reduced growth factor Matrigel diluted in M87A medium at 2% v/v, (0.18 – 0.2 mg/ml) at room temperature for at least 2 hr. The coverslips were washed once with M87A medium prior to seeding FACS-sorted LEP and MEP at 5,000 cells per 0.7 cm$^2$ chamber. After allowing initial attachment in the incubator for 1 h, the coverslips were washed again, and fixed in 4% PFA after 4 h. The samples were stained for K14 and K19 to confirm cell identity, and phalloidin was used to label cell cortex. Z-stacks of healthy, radially symmetric cells not touching nearby cells were acquired using a 63x/NA 1.4 oil objective at 0.26 $\mu$m z- resolution on a Zeiss Cell Observer Z1, Yokagawa spinning disk microscope. The FIJI angle tool was used to measure the contact angle at the cell-ECM interface at four points around the circumference of each cell, taken from coronal and sagittal projections. The gross cell shape was used for measurements and any thin, flat membrane extrusions were discarded. The four measurements were averaged on a per-cell basis for analysis.

**1.5.3 Cell-cell contact angles:** Cell-cell contact angle measurements were described previously[2]. FACS-sorted GFP+ LEP and mCherry+ MEP were centrifuged into small oblong agarose microwells designed to hold two cells (20 $\mu$m x 40 $\mu$m x 20 $\mu$m) and cultured for 3 h. Healthy pairs of similar-sized cells were imaged using a 40X/NA 1.1 water objective using a Zeiss Cell Observer Z1, Yokagawa spinning disk microscope. The FIJI angle tool was used to measure four contact angles at each cell-cell interface, which were averaged for each cell doublet for analysis.

**1.5.4 Interfacial tension calculations:** Young's equation was used to calculate the average interfacial tension of the various cell-cell and cell-ECM interfaces based on the cortical tension and contact angle measurements[2]. The cell-cell interfacial tension ($\gamma_{\text{cc}}$) was calculated as

$$\gamma_{\text{cc}} = \gamma_{\text{c}} \cos \theta_{\text{cc}} , \quad (5)$$

where $\gamma_{\text{c}}$ is the cell cortical tension and $\theta_{\text{cc}}$ is the average cell-cell contact angle.

The cell-ECM interfacial tension ($\gamma_{\text{cx}}$) was calculated as

$$\gamma_{\text{cx}} = \gamma_{\text{c}} \cos \theta_{\text{cx}} + \gamma_x , \quad (6)$$

where $\gamma_{\text{c}}$ is the cell cortical tension, $\theta_{\text{cx}}$ is the average cell-ECM contact angle, and $\gamma_x$ is the

ECM-medium tension. As only relative differences in LEP- and MEP- ECM tensions are relevant, we assigned $\gamma_{\text{MEP-ECM}}$ (the lowest tension interface) to be 0.

To get confidence intervals for the interfacial tensions, we calculated their variance using error propagation as

$$\sigma^2_{\gamma_{\text{cc}}} = \gamma^2_{\text{cc}} \left( \frac{\sigma^2_{\gamma_{\text{c}}}}{\gamma^2_{\text{c}}} + \sigma^2_{\theta_{\text{cc}}} \tan^2 \theta_{\text{cc}} \right) \text{, and} \tag{7}$$

$$\sigma^2_{\gamma_{\text{cx}}} = \gamma^2_{\text{cx}} \left( \frac{\sigma^2_{\gamma_{\text{c}}}}{\gamma^2_{\text{c}}} + \sigma^2_{\theta_{\text{cx}}} \tan^2 \theta_{\text{cx}} \right) . \tag{8}$$

#### 1.6 Cell motility analysis

To measure undirected cell motility in the absence of cell sorting, we tracked nuclei in spheroids containing a single cell type (for e.g., MEP or LEP only spheroids). Cell cytoplasms were labeled with cytoplasmic GFP and roughly 10% of the nuclei were additionally labeled with H2B-mScarlet. Spheroids containing 2-10 labeled nuclei, with no neighboring organoids and diameter between 60 and 120 $\mu\text{m}$ were chosen for time lapse imaging. For each organoid, seven z-slices 5 $\mu\text{m}$ apart were acquired every 10 minutes.

**1.6.1 Cell tracking:** Nuclei were tracked using the Trackmate plugin[7] in FIJI. Only tracks longer than 5 h with less than two gaps were selected. Cell positions were corrected relative to the centroid of the spheroid to account for organoid translation. Only the cell displacements in $x$-and $y$-directions were used due to the high uncertainty in the $z$-position (5 $\mu\text{m}$ z-resolution). Instantaneous cell speeds were measured as the distance a cell moved per frame as

$$v_i = \frac{\mathbf{r}_i - \mathbf{r}_{i-1}}{t_i - t_{i-1}} , \tag{9}$$

where $\mathbf{r_i}$ is the position at time $t_i$.

**1.6.2 Cell diffusion analysis:** Single cell motility was coupled to whole organoid translation and rotation. This was especially pronounced for organoids in agarose, where the relative magnitudes of cell displacement and organoid motion were comparable. To overcome this, we instead measured the change in distance between a pair of nuclei (**Fig. 4B**). The mean squared relative displacement (MSRD) between nuclei was calculated by averaging all measurements within an organoid as

$$MSRD_{\Delta t} = \overline{(r_{t+\Delta t} - r_t)^2} . \tag{10}$$

To elaborate on the nature of cell dynamics, we further examined the relationship between MSRD and lag times (**Sup. Fig. 4A**). We checked for anomalous diffusion by fitting data to the equation

$$MSRD = 2\, D_{\text{eff}}\, \Delta t^{\alpha}, \tag{11}$$

where $D_{\text{eff}}$ and $\alpha$ are the effective diffusion coefficient and exponents. Only lag times less than 3 h were considered to minimize the confounding effects from movement in a confined volume. The average value of $\alpha$ was 1.1 indicating near-diffusive cell dynamics (**Sup. Fig. 4B**). Therefore, we used a simple diffusion model ($\alpha = 1$) to estimate $D_{\text{eff}}$.

#### 1.7 Immunofluorescence

**1.7.1 2D immunofluorescence:** Cells were fixed in 4% PFA for 10 min at room temperature and permeabilized with 0.5% Triton X-100 for 5 min at 4 $^{\circ}\mathrm{C}$. The samples were blocked for 1 h at room temperature in blocking buffer (PBS + 10% heat-inactivated goat serum + 0.1% w/v BSA + 0.2% v/v Triton X-100 + 0.04% v/v Tween-20) and incubated overnight at 4 $^{\circ}\mathrm{C}$ with primary antibodies solution prepared in the blocking buffer. Next, the samples were rinsed in wash buffer (PBS + 0.1% w/v BSA + 0.2% v/v Triton X-100 + 0.04% v/v Tween-20) 3 times for 5 min at room temperature, incubated with secondary antibody in blocking buffer for 30 min at room temperature, rinsed in wash buffer 4 times for 5 min at room temperature, and incubated with DAPI or phalloidin in PBS for 10 min in room temperature. The details for all antibodies used provided in **Table 3**.

**1.7.2 3D immunofluorescence:** Cells were fixed in pre-warmed 2% PFA for 45 min at room temperature, permeabilized with 0.5% Triton X-100 for 15 min at room temperature, blocked with blocking buffer for 2 h at room temperature or overnight at 4 $^{\circ}\mathrm{C}$. If using mouse-origin primary antibodies, samples were also incubated overnight with 1:50 dilution mouse Fab fragment (Jackson ImmunoResearch #115-007-003). Samples were then incubated with primary antibody in blocking buffer for 24-48 h at 4 $^{\circ}\mathrm{C}$, rinsed in wash buffer 3 times for 1 h at room temperature, incubated with secondary antibody in blocking buffer overnight at 4 $^{\circ}\mathrm{C}$, and rinsed in wash buffer 3 times for 1 hr at room temperature. Samples were stored in PBS at 4 $^{\circ}\mathrm{C}$ prior to imaging. The details for all antibodies used provided in **Table 3**.

#### 1.8 Graphing and statistical analysis

All data compilation, graphing and statistical analysis was done with R using RStudio. Details for sample numbers and replicates is outlined in the figure legends. Each experimental replicate was validated against the existing data set to ensure exclusion of technical outliers (for example due to cell culture stress). Two-tailed non-parametric Wilcoxon tests were used for comparing two groups unless stated otherwise. The Holm-Bonferroni test was used to adjust p-values when comparing multiple groups. A priori power analysis was used to determine minimum sample sizes. For example, to detect a 10% shift in $\phi_{\text{b}}$ with $\sigma = 0.14$ we targeted a minimum sample size of 50 for power=0.95. Bootstrapping was used to identify confidence intervals for model fitting of experimental data where applicable. The goodness of fit was evaluated by the examination of residuals.

### 2 Computational lattice model for mammary organoids

#### 2.1 2D hexagonal lattice description

Tissue were modeled as a 2D hexagonal lattice with the radius equal to $R$ cell lengths (**Supp. Fig. 3A**). Each cell was defined as a regular hexagon with edge length $a$ equal to 10 $\mu\mathrm{m}$. The contact length between any two neighbors was $a$ $\mu\mathrm{m}$, and corresponding interface area was $10a$ $\mu\mathrm{m}$. The cell centers were $2a$ and $\sqrt{3}a$ apart in the $x$- and $y$-dimensions respectively. Lattice points less than $R * a$ away from the tissue center were assigned as *cells* and points between $R * a$ and $(R + 1) * a$ from the tissue center were assigned as *ECM*. Any cells with 1 or more ECM neighbors was labeled a *boundary* cell, and all others were *core* cells. Unless specified otherwise, simulated tissues had a diameter 140 $\mu\mathrm{m}$, corresponding to 7 cells along the tissue diameter. For this geometry, a tissue had 53 cells, of which 24 were boundary cells.

#### 2.2 3D body-centered cubic lattice description

Tissues were modeled as a 3D body-centered cubic (BCC) lattice, a uniform polyhedral foam of near-minimal surface area (**Supp. Fig. 3B**). In this model, cells are considered truncated octahedrons with edge length $a$ $\mu\mathrm{m}$. Each cell has 6 nearest neighbors (square interface) and 8 next-nearest neighbors (hexagonal interface). The tissue surface was considered a sphere of equivalent diameter to avoid overestimating the tissue surface area in contact with the ECM. Cell centers are uniformly spaced in $x$-, $y$- and $z$-dimensions by $2(\sqrt{2} - 1) * a$ $\mu\mathrm{m}$. Lattice points less than $R * a$ away from the tissue center were assigned as *Cells* and points between $R * a$ and $(R + 1) * a$ from the tissue center were assigned as *ECM*. Boundary cells were defined as cells that had >1 nearest neighbors as ECM. Unless specified otherwise, simulated tissues had a diameter 130 $\mu\mathrm{m}$, corresponding to 9 cells along the tissue diameter. For this geometry, a tissue had 281 cells, of which 134 were boundary cells.

#### 2.3 Calculations for macrostate degeneracy

Each cell site within the simulated tissue was assigned an identity (1: LEP and 2: MEP) either randomly or by selecting a target boundary LEP occupancy ($\phi_\mathrm{b}$). This process was repeated multiple times to get an ensemble of tissue configurations (microstates). For the estimation of macrostate degeneracy, $10^5$ tissue configurations were sampled randomly to examine the distribution of $\phi_\mathrm{b}$. For a target $\phi_\mathrm{b}$ (macrostate), the number of LEP in boundary and core were held constant, and LEP position was sampled randomly in each compartment separately. Any configurations that would require the number of LEP in the boundary or core to exceed the total number of LEP were excluded from analysis. For each $\phi_\mathrm{b}$, $10^4$ configurations were generated randomly.

##### 2.4 Calculations for the average mechanical energy

There are 5 types of interfaces in the tissue - 2 cell-ECM interfaces (LEP-ECM and MEP-ECM) and 3 cell-cell interfaces (LEP-LEP, LEP-MEP and MEP-MEP). The tensions for each interface was measured experimentally (refer to section **??**, **Fig. 3B-E**). The total area corresponding to each type of interface was calculated by summation of surface areas across all tissue interfaces. The mechanical energy of a tissue configuration was defined as the sum of the product of interfacial tensions and total areas for each type of the interface (**Fig. 3A**).

The mechanical energy was largely determined by the fraction of LEP-ECM contacts ($\phi_{\text{b}}$)(**Fig. 3G**). While different microstates with the same $\phi_{\text{b}}$ had slightly different energies, the energies were symmetric about the average value. Further, these differences were smaller than the energy difference across macrostates (**Supp. Fig. 3D**). The number of LEP-LEP or LEP-MEP contacts were poor predictors of energy (**Supp. Fig. 3C**). We defined the mechanical potential ($\Delta$E) as the slope of the average mechanical energy against $\phi_{\text{b}}$. To get confidence intervals for $\Delta$E, we used bootstrapping (n=1000) to sample 1000 configurations for each $\phi_{\text{b}}$, averaged their energies and performed a linear fit (**Supp. Fig. 3E**).

#### 3 Analytical lattice model for mammary organoids

##### 3.1 Lattice description and structural metrics

We built a lattice-based model, where organoids were modeled as spheres with diameter $d$ with two compartments - *core* and *boundary* (**Supp. Fig. 3F**). The thickness of the boundary layer was set as $t$. All cells were assumed to have the same volume ($v_{\text{c}}$) and the tissue had no interstitial space. Therefore, the total number of lattice sites (*cells*) $N_t$ was

$$N_{\text{t}} = \frac{1}{6}\pi d^3/v_{\text{c}} \; . \tag{12}$$

The number of cells in each compartment ($N_{\text{c}}$ and $N_{\text{b}}$) was estimated by dividing the compartment volume by $v_{\text{c}}$ as

$$N_{\text{c}} = \frac{4}{3}\pi(d/2 - t)^3/v_{\text{c}} \; , \tag{13}$$

and

$$N_{\text{b}} = N_{\text{t}} - N_{\text{c}} \; . \tag{14}$$

The fraction of sites in the tissue boundary, $F_{\text{b}}$ was

$$F_{\text{b}} = N_{\text{b}}/N_{\text{t}} \; . \tag{15}$$

Based on volume conservation, $v_{\text{c}}$ was set to be equal to the volume of a sphere of diameter 20 $\mu\text{m}$, the size for single HMEC in suspension. Therefore, an 80 $\mu\text{m}$ diameter tissue was estimated to have

64 cells, of which half were in the boundary compartment (**Supp. Fig. 3H**).

Organoids contained two types of cells - LEP and MEP. The total fraction of LEP in the tissue was $\Phi$, and was held constant. We used the fraction of LEP in the boundary and core ($\phi_{\mathsf{b}}$ and $\phi_{\mathsf{c}}$ respectively) as descriptors of tissue structure. This framework can be easily generalized to other tissue geometries or to incorporate additional cell types. The numbers of total, boundary and core LEP ($K_{\mathsf{t}}$, $K_{\mathsf{b}}$, and $K_{\mathsf{c}}$ respectively) were

$$K_{\mathsf{t}} = \Phi \; N_t \;, \tag{16}$$

$$K_{\mathsf{b}} = \phi_{\mathsf{b}} \; N_{\mathsf{b}}, \text{ and} \tag{17}$$

$$K_{\mathsf{c}} = \phi_{\mathsf{c}} \; N_{\mathsf{c}} = K_{\mathsf{t}} - K_{\mathsf{b}}. \tag{18}$$

As the total number of LEP was fixed, $\phi_{\mathsf{c}}$ was replaced as a function of $\phi_{\mathsf{b}}$ :

$$\phi_{\mathsf{c}} = \frac{\Phi - \phi_{\mathsf{b}} \; F_{\mathsf{b}}}{1 - F_{\mathsf{b}}} \;. \tag{19}$$

All possible values of $\phi_{\mathsf{b}}$ might not be possible due to constraints on the available number of LEP and the distribution of sites between the boundary and core compartments. For example, the number of LEP in the boundary ($K_{\mathsf{b}}$) or core ($K_{\mathsf{c}}$) cannot exceed the total number of LEP ($K_t$). The maximum and minimum number of LEP in the boundary were estimated as

$$K_{\mathsf{b,\,max}} = \mathsf{min}(K_t;\; N_{\mathsf{b}}), \text{ and} \tag{20}$$

$$K_{\mathsf{b,\,min}} = \mathsf{max}(0;\; K_t - N_{\mathsf{c}}). \tag{21}$$

All values of $\phi_{\mathsf{b}}$ that do not follow $K_{\mathsf{b,\,min}} < K_{\mathsf{b}} < K_{\mathsf{b,\,max}}$ were assigned the probability $0$.

#### 3.2 Calculations for macrostate degeneracy

The macrostate degeneracy ($\Omega$) represents the number of microstates ($m$) corresponding to a particular macrostate ($M$), $\phi_{\mathsf{b}}$ (**Supp. Fig. 3G**). We used combinatorics to calculate the number of ways to distribute $K_{\mathsf{b}}$ and $K_{\mathsf{c}}$ LEP across $N_{\mathsf{b}}$ boundary and $N_{\mathsf{c}}$ core sites respectively as

$$\Omega_b = \binom{N_{\mathsf{b}}}{K_{\mathsf{b}}} \quad \text{and} \quad \Omega_c = \binom{N_{\mathsf{c}}}{K_{\mathsf{c}}}. \tag{22}$$

Therefore, the degeneracy of the $i^{th}$ macrostate (**Fig. 3H**) was calculated as

$$\Omega_i = \Omega_{b,i} \;.\; \Omega_{c,i} = \binom{N_{b,i}}{K_{b,i}} \binom{N_{c,i}}{K_{c,i}}. \tag{23}$$

In the absence of any driving forces, all microstates had equal probability ($p$), and the sum of all microstate probabilities added up to 1. Therefore,

$$\sum_m p = \sum_M \Omega_i \, p = 1. \tag{24}$$

Consequently, the macrostate probabilities under these conditions were

$$p_{M,i} = \frac{\Omega_i}{\sum_M \Omega_i} \, . \tag{25}$$

Unsurprisingly, this distribution takes the shape of a normal distribution which is also a maximum entropy distribution for a fixed mean and standard deviation. The mean for this distribution was $\Phi$, and its width was determined by tissue geometry ($d$ and $F_\text{b}$), where $d$ determined the total number of available sites and $F_\text{b}$ determined the distribution of sites between the boundary and core compartments. The degeneracy-based probability distribution and the measured distribution of MEP spheroids had identical means, but the experiments had higher variance. This increased variance was due to hidden degrees of freedom along other structural metrics and the technical uncertainty in our measurements, which we additionally incorporated into the analytical model (see section **??**).

A similar approach can be also used to estimate the entropy-based distributions for non-spherical tissues (e.g. tubes or tissues with lumens) or tissues with different constraints on cell deformability or connectivity. The values of $N_t$, $N_\text{b}$ and $N_\text{c}$ will vary with the geometric constraints of the system. We also verified these relationships using 2D and 3D lattice models (see section **??**).

##### 3.3 Calculations for average mechanical energy

The mechanical energy of a tissue arises from the surface tensions (and energy) at the various cell-cell and cell-ECM interfaces. These tensions are the combined effects of cell adhesion (via molecular binding interactions of adhesion molecules on the outer surfaces of the cell membranes) and cell contractility (via the active contractions of the actomyosin cytoskeleton coupled to these adhesion molecules). The critical role for interfacial tensions in determining cell sorting is well established in multiple tissue systems[8,9]. In the mammary gland, there are three types of cell-cell interfaces (LEP-MEP, LEP-MEP and MEP-MEP) and two types of cell-ECM (LEP-ECM and MEP-ECM) interfaces. Each type of interface had a unique tension ($\gamma$), and all five interfacial tensions were quantified experimentally (Section **??**). The mechanical energy of each cell interface is the product of its tension and surface area, and the total mechanical energy of a tissue microstate is the sum of the mechanical energies across all its interfaces. While different microstates within a macrostate likely have different mechanical energies, each macrostate can still be represented by an average mechanical energy using a mean field approximation[10]. This approach is valid for energies symmetrically distributed about the average energy, which we additionally confirmed using computational methods (**Supp. Fig. 3D**).

In the lattice model with a boundary and core compartment, the cell interfaces can be between two core cells, between a core and a boundary cell, between two boundary cells, and between a boundary cell and the ECM. Therefore, the total mechanical energy was

$$E_{\text{total}} = E_{\text{core-core}} + E_{\text{core-boundary}} + E_{\text{boundary-boundary}} + E_{\text{boundary-ECM}} \ . \tag{26}$$

We used mean field approximation to estimate each of the above terms based on the probability of each type of cell interface (LEP-ECM, MEP-ECM, LEP-LEP, LEP-MEP, and MEP-MEP) within different tissue compartments. For example, if $\phi_{\text{c}}$ was the LEP core occupancy, the probabilities of a core lattice site being occupied by a LEP or a MEP were $\phi_{\text{c}}$ and $(1-\phi_{\text{c}})$ respectively. Consequently, the probability of LEP-MEP interactions in the tissue core was

$$p^{\text{LM}}_{\text{core}} = p^{\text{LEP}}_{\text{core}} \cdot p^{\text{MEP}}_{\text{core}} = \phi_{\text{c}} \ . \ (1-\phi_{\text{c}}) \ . \tag{27}$$

Therefore, the average interfacial energy for core-core interactions was

$$\begin{aligned} E_{\text{core-core}} &= A_{\text{cc}}[p^{\text{LL}}_{\text{core}}\ \gamma_{\text{ll}} + p^{\text{LM}}_{\text{core}}\ \gamma_{\text{lm}} + p^{\text{MM}}_{\text{core}}\ \gamma_{\text{mm}}] \\ &= A_{\text{cc}}\left[\phi_{\text{c}}^2\gamma_{\text{ll}} + 2\phi_{\text{c}}(1-\phi_{\text{c}})\gamma_{\text{lm}} + (1-\phi_{\text{c}})^2\gamma_{\text{mm}}\right] \\ &= A_{\text{cc}}\left[\phi_{\text{c}}^2(\gamma_{\text{ll}} - 2\gamma_{\text{lm}} + \gamma_{\text{mm}}) + 2\phi_{\text{c}}(\gamma_{\text{lm}} - \gamma_{\text{mm}})\right] \\ &= A_{\text{cc}}\left[\phi_{\text{c}}^2 E_1 + \phi_{\text{c}} E_2\right], \end{aligned} \tag{28}$$

where $A_{\text{cc}}$ was the total surface area for core-core interactions, $E_1 = \gamma_{\text{ll}} - 2\gamma_{\text{lm}} + \gamma_{\text{mm}}$, and $E_2 = 2(\gamma_{\text{lm}} - \gamma_{\text{mm}})$.

Similarly, we calculated the other terms in the Equation **??** as

$$\begin{aligned} E_{\text{core-boundary}} &= A_{\text{bc}}\left[\phi_{\text{c}}\phi_{\text{b}}\gamma_{\text{ll}} + \{\phi_{\text{c}}(1-\phi_{\text{b}}) + \phi_{\text{b}}(1-\phi_{\text{c}})\}\gamma_{\text{lm}} + (1-\phi_{\text{c}})(1-\phi_{\text{b}})\gamma_{\text{mm}}\right] \\ &= A_{\text{bc}}\left[\phi_{\text{c}}\phi_{\text{b}}(\gamma_{\text{ll}} - 2\gamma_{\text{lm}} + \gamma_{\text{mm}}) + (\phi_{\text{c}} + \phi_{\text{b}})(\gamma_{\text{lm}} - \gamma_{\text{mm}})\right] \\ &= A_{\text{bc}}\left[\phi_{\text{c}}\phi_{\text{b}}E_1 + \frac{(\phi_{\text{c}} + \phi_{\text{b}})}{2}E_2\right] \ , \end{aligned} \tag{29}$$

.

$$\begin{aligned} E_{\text{boundary-boundary}} &= A_{\text{bb}}\left[\phi_{\text{b}}^2\gamma_{\text{ll}} + 2\phi_{\text{b}}(1-\phi_{\text{b}})\gamma_{\text{lm}} + (1-\phi_{\text{b}})^2\gamma_{\text{mm}}\right] \\ &= A_{\text{bb}}\left[\phi_{\text{b}}^2(\gamma_{\text{ll}} - 2\gamma_{\text{lm}} + \gamma_{\text{mm}}) + 2\phi_{\text{b}}(\gamma_{\text{lm}} - \gamma_{\text{mm}})\right] \\ &= A_{\text{bb}}\left[\phi_{\text{b}}^2 E_1 + \phi_{\text{b}} E_2\right] \ \text{, and} \end{aligned} \tag{30}$$

$$\begin{aligned} E_{\text{boundary-ECM}} &= A_{\text{x}}\left[\phi_{\text{b}}\gamma_{\text{lx}} + (1-\phi_{\text{b}})\gamma_{\text{mm}}\right] \\ &= A_{\text{x}}\left[\phi_{\text{b}}E_{\text{x}} + \gamma_{\text{mm}}\right], \end{aligned} \tag{31}$$

where $E_{\text{x}} = \gamma_{\text{lx}} - \gamma_{\text{mx}}$. From the measured interfacial tensions (see section **??**), $\gamma_{\text{lm}}$ was roughly the average of the two homotypic tensions ($\gamma_{\text{ll}}$ and $\gamma_{\text{mm}}$), implying $E_1 \approx 0$. Therefore, the total mechanical energy (Eq **??**) was simplified to a linear function of $\phi_{\text{b}}$ as

$$E_{\text{total}} = [A_{\text{cc}}\phi_{\text{c}} + A_{\text{bc}}\frac{(\phi_{\text{b}} + \phi_{\text{c}})}{2} + A_{\text{bb}}\phi_{\text{b}}]E_2 + A_{\text{x}}E_{\text{x}}\phi_{\text{b}} \\ = \Delta E\phi_{\text{b}} + C. \tag{32}$$

The linear scaling of mechanical energy with $\phi_{\text{b}}$ was consistent with the energy function predicted for the maximum entropy distribution with a given average $\phi_{\text{b}}$ (Eq. **??**), with $\beta = \frac{\Delta E}{a}$ where ΔE was the mechanical potential and $a$ was the tissue activity. $\Delta$E can be numerically calculated for different cell geometries and connectivities. Here, we used a body-centered cubic (BCC) lattice to model computationally calculate the interfacial areas and $\Delta$E (see section **??**). For smaller tissues, the large difference in cell-ECM tensions ($E_{\text{x}}$) dominated the mechanical potential. However, for larger tissues with lower surface area to volume ratio, the cell-cell interactions have a higher weight. Additionally, the mean field approximation only holds if the structural energy is solely a function of $\phi_{\text{b}}$, which is the case for mammary organoids $< 150$ μm in diameter. In much larger tissues, the differences in cell-cell adhesions can lead to the formation of sub-domains of a single cell type. In such cases, the resulting structures are not well captured by a single structural metric and the energy function must be modified accordingly.

### 4 Modeling organoid ensembles as maximum entropy distributions

#### 4.1 Maximum entropy distributions constrained by a defined average tissue structure

Mammary organoids had highly reproducible structural averages and variances across experimental and biological replicates(**Supp. Fig. 1L,M**). Statistical mechanics offers a powerful framework for the quantitative description of these reproducible distributions, where the distribution which maximizes the system entropy under known constraints is the most likely distribution. In thermodynamic systems, these constraints primarily arise from a constant average mechanical energy of particles. Without prior knowledge of the source of mechanical energy in organoid ensembles, we began by estimating the maximum entropy distribution for a fixed ensemble average structure using an information theoretic approach. We assumed that the total number of cells remained fixed. Here, we used the LEP boundary occupancy, $\phi_{\text{b}}$ as the descriptor of tissue structure, but this approach can also be generalized to other metrics of tissue structure.

Along this single structural descriptor, a large number of tissue configurations can have the same $\phi_{\text{b}}$ arising from additional degrees of freedom along other structural features. Therefore, we defined all possible tissue configurations as a **microstate (m)**, and the collection of all configurations with a given $\phi_{\text{b}}$ the **macrostate (M)**. For simplicity, we divided tissues into 10 measurable macrostates ($M_i$) with bins centered at $\phi_{b,i} \in (0.05, 0.15, \,...\,, 0.95)$. The macrostate degeneracy ($\Omega_i$) is defined as the number of microstates in $M_i$ (or $\phi_{\text{b}} = \phi_{b,i}$). Assuming all microstates within a macrostate are equally probable (microstate probability = $p_i$), the macrostate probability was defined as:

$$p_{M,\,i} = \sum_{m \in M_i} p_i = \Omega_i \; p_i \; . \tag{33}$$

The Shannon entropy for a given probability distribution is

$$S = -\sum_{m} p_i \; ln(p_i\;) = \sum_{M} p_{M,\,i} \; ln(p_i\;) = \sum_{M} \Omega_i \; p_i \; ln(p_i\;). \tag{34}$$

Prior to maximizing the entropy function, we applied the following constraints to the distributions:

- The sum of all microstate probabilities was 1

$$\sum_{m} p_i = \sum_{M} p_{M,\,i} = \sum_{M} \Omega_i \; p_i = 1\;. \tag{35}$$

- The average boundary LEP occupancy was $\overline{\phi_{\text{b}}}$

$$\sum_{m} p_i.\phi_{b,i} = \sum_{M} \Omega_i \; p_i \; \phi_{b,i} = \overline{\phi_{\text{b}}}\;. \tag{36}$$

Lagrange multipliers are commonly used to find minima or maxima of a functions subject system constraints. Therefore, we defined an augmented entropy function that combined the Shannon entropy with the constraints for normalization and an average observed structure using the Lagrange multipliers $\lambda$ and $\beta$ respectively.

$$S' = -\sum_{M} \Omega_i \; p_i \; ln(p_i) - \lambda \left[\sum_{M} \Omega_i \; p_i - 1\right] - \beta \left[\sum_{M} \Omega_i \; p_i \; \phi_{b,i} - \overline{\phi_{\text{b}}}\right]\;. \tag{37}$$

Maximizing $S'$ with respect to $p_i$, we get

$$\frac{\partial S'}{\partial p_i} = 0 = -ln(p_i) - 1 - \lambda - \beta\phi_{b,i}\;. \tag{38}$$

Therefore, the macrostate probability is

$$p_{M,\,i} = \Omega_i \; p_i \; = \Omega_i \; e^{-1-\lambda}\;.\; e^{-\beta\phi_{b,i}} = \frac{\Omega_i}{Z}\;.\; e^{-\beta\phi_{b,i}}\;. \tag{39}$$

where $Z$ is the partition function

$$Z = \sum_{M} \Omega_i \; e^{-\beta\phi_{\text{b}}}\;. \tag{40}$$

In Boltzmann statistics, the probability of occupancy of a macrostate is a function of its potential energy and degeneracy, and is described as

$$p_i = \frac{\Omega_i}{Z}\;.\; e^{-\frac{E_i}{k_B T}}\;, \tag{41}$$

where $E_i$ is the average potential energy and $k_BT$ is a scaling factor that corresponds to energy required for unit increase in entropy. Analogously, in our system, $\beta\phi_{b,i}$ should also represent a dimensionless mechanical energy. We hypothesize that this mechanical energy stems from the surface interactions at the cellular interfaces, and we calculated it both analytically (section **??**) or computationally (section **??**). In the absence of driving forces $\beta$=0 and $p_{M,\,i} = \Omega_i/Z$, implying equal probability for all possible microstates. Therefore, we used the distribution of MEP spheroids to experimentally measure macrostate degeneracy as all cells in these tissues are identical. We expect $\Omega_i$ to only be a function of cell and tissue geometry. To confirm this, we additionally made spheroids containing only LEP, MCF10A or MCF7 cells, which all had similar distributions (**Supp. Fig. 3M**).

##### 4.2 Calculation for the maximum entropy distribution

We assumed the tissue exchanges the energy, but not cells with the system. Therefore, we modeled organoids at steady state as a canonical ensemble, where their structural distribution can described as a maximum entropy distribution (Eq. **??**). The macrostate degeneracy ($\Omega_i$) was was either calculated analytically (Eq. **??**) or experimentally (from the distribution of MEP spheroids with similar size and composition). The mechanical potential ($\Delta$E) was calculated analytically (Eq. **??**) or computationally (Section **??**). Tissue activity was estimated by fitting experimental distributions to Eq. **??** with calculated values of $\Omega_i$ and $\Delta$E, where activity = $\Delta E/\beta$. Bootstrapping was used to estimate the confidence intervals for activity, where 100 organoids were sampled with replacement 1000 times.

Tissue activity cannot be determined analytically as it is an emergent property of tissues, and arises from cellular fluctuations. However, as it represents the kinetic component of energy, we examined the correlation between the inferred activity from the model and the measured cellular diffusion coefficients across various conditions.

##### 4.3 Technical sources for structural variance

The analytical calculations of structural degeneracy using combinatorics (Eq. **??**) resulted in structural distributions with similar averages but larger variance compared to experimentally measured distribution of organoids. We hypothesized that this greater variance could be attributed to additional degrees of freedom along different structural metrics (e.g., deviations in cell shape) or technical uncertainty in our measurements. A major source of technical uncertainty could stem from the analysis of only the middle three sections of the organoids, resulting in variance from sampling of a subset of cells within the organoid. A hypergeometric distribution describes the probability of success when drawing a sample without replacement, and enabled the quantification of technical variance in the measurements. Therefore, we used this approach to analytically calculate the average and variance for the measured LEP fraction ($\Phi^m$) and LEP boundary occupancy ($\phi_{\mathsf{b}}^m$).

For an imaging thickness of $2s$ (**Supp. Fig. 3I**), we estimated the number of sites in the sample volume in each compartment based on the compartment volume as

$$n_{\mathsf{t}} = 2\pi s(R^2 - s^2)/v_{\mathsf{c}}, \tag{42}$$

$$n_{\mathsf{c}} = min\{2\pi s((R-t)^2 - s^2)/v_{\mathsf{c}}\ ;\ \frac{4}{3}\pi(R-t)^3\}, \tag{43}$$

and

$$n_{\mathsf{b}} = n_{\mathsf{t}} - n_{\mathsf{c}}. \tag{44}$$

The number of LEP in each tissue compartment within the sampled volume was

$$\text{Total LEP, } k_\text{t} = \Phi^m \;.\; n_\text{t}, \tag{45}$$

$$\text{Boundary LEP, } k_\text{b} = \phi_\text{b}^m \;.\; n_\text{b}, \text{ and} \tag{46}$$

$$\text{Core LEP, } k_\text{c} = k_\text{t} - k_\text{b}. \tag{47}$$

The fraction of boundary sites in the sample volume, $f_\text{b}$, was

$$f_\text{b} = \frac{n_\text{b}}{n_\text{t}}. \tag{48}$$

As expected, $f_\text{b}$ was smaller than $F_\text{b}$, implying a higher sampling rate for core sites in the mammary organoids. The $F_\text{b}$ and $f_\text{b}$ converged for imaging depths comparable to the tissue diameter. We separately analyzed the LEP distribution in the boundary and core to account for any discrepancy between the measured and true $\Phi$ and $\phi_\text{b}$ by modeling the sampling process as a hypergeometric distribution. The sample and population averages are expected to be the same for hypergeometric sampling. Therefore, we expect the average for measured boundary and core LEP occupancy to be the same as the true LEP occupancies, and were accordingly defined as

$$\overline{\phi_\text{b}^m} = \frac{k_\text{b}}{n_\text{b}} = \frac{K_\text{b}}{N_\text{b}} = \phi_\text{b}. \tag{49}$$

Similarly, the average core LEP fraction in the sample volume is

$$\overline{\phi_\text{c}^m} = \frac{k_\text{c}}{n_\text{c}} = \frac{K_\text{c}}{N_\text{c}} = \phi_\text{c}. \tag{50}$$

However, due to increased sampling of core sites ($f_\text{b} > F_\text{b}$), we predicted a shift in $\Phi^m$ compared to $\Phi$. This relationship was derived as

$$\begin{aligned}
\Phi^m &= \frac{k_\text{b} + k_\text{c}}{n_\text{t}} \\
&= \phi_\text{b}\frac{n_\text{b}}{n_\text{t}} + \phi_\text{c}\left(1 - \frac{n_\text{b}}{n_\text{t}}\right) \\
&= \phi_\text{b}\frac{n_\text{b}}{n_\text{t}} + \frac{\Phi - \phi_\text{b}.F_\text{b}}{1 - F_\text{b}}\left(1 - \frac{n_\text{b}}{n_\text{t}}\right) \\
&= \phi_\text{b}\frac{f_\text{b} - F_\text{b}}{1 - F_\text{b}} + \Phi\frac{1 - f_\text{b}}{1 - F_\text{b}}.
\end{aligned} \tag{51}$$

As a result, for sorted mammary organoids that have more LEP in the tissue core, $\Phi^m$ was expected to be higher than $\Phi$ (**Supp. Fig. 3J**). We corrected for this difference in the measured and estimated LEP fraction for the organoids prior to fitting the experimental data to the analytical model. An additional consequence of volume sampling was increased variance in the measurements, which was estimated as

$$\text{var}(\phi_{\text{b}}^m) = \phi_{\text{b}}(1-\phi_{\text{b}})\frac{N_{\text{b}}-n_{\text{b}}}{N_{\text{b}}-1}.\frac{1}{n_{\text{b}}} \ . \tag{52}$$

For tissues containing equal proportion of LEP and MEP, the variance is highest at $\phi_{\text{b}} = 0.5$ (**Supp. Fig. 3K**). This variance decreased as we image more sites ($n_{\text{b}}$), either by increasing sampling ratio ($n_{\text{t}}/N_{\text{t}}$) or tissue size ($d$). For our setup, we expect this variance to be less than 0.02. We incorporated the var($\phi_{\text{b}}^m$) by applying Gaussian filters to the original probability function. This had a widening effect on the predicted distributions (**Supp. Fig. 3L**). The resulting analytical predictions accurately matched the variance of the experimental data, while the technical variance or entropy alone would predict much narrower distributions (**Supp. Fig. 3N**).

#### 4.4 Favorability of LEP partitioning between the tissue boundary and core

The tissue state variables (average mechanical energy, degeneracy and activity) determine the distribution of tissue ensembles, thereby controlling the balance between structural order and disorder. The extent of structural order is regulated by the relative forward and reverse rates of LEP translocation from the tissue core to boundary ($k_{\,c\to b}$ and $k_{\,b\to c}$). The average occupancy of LEP in the boundary and core compartments was calculated from the analytical model, from which an effective equilibrium constant ($K_{\text{eq}}$) for cell translocation can be estimated as

$$\begin{aligned} K_{\text{eq}} &= \frac{k_{\,c\to b}}{k_{\,b\to c}} = \frac{\text{[bdr. LEP] [core MEP]}}{\text{[core LEP] [bdr MEP]}} \\ &= \frac{\phi_{\text{b}}\,(1-\phi_{\text{c}})}{\phi_{\text{c}}\,(1-\phi_{\text{b}})}. \end{aligned} \tag{53}$$

Using analogy to equilibrium systems, the free energy change for cell translocation ($\Delta$G), which determines the favorability of this process, is proportional to -log($K_{\text{eq}}$). A negative $\Delta$G implies LEP have a tendency to be enriched in the tissue boundary, while a positive $\Delta$G implies LEP exclusion from the boundary.

For validation of experimental results, the average $\phi_{\text{b}}$ was used to estimate $\phi_{\text{c}}$, assuming $F_{\text{b}} = 0$. We used bootstrapping to get the average and confidence intervals for $\Delta$G for each condition, and compared results to the model predictions.

#### Supplemental Tables

Table 1: **List of abbreviations.**

| Abbreviation | Description |
|---|---|
| $\Phi$ | LEP or GFP fraction. |
| $\phi_{\text{b}}$ | LEP or GFP fraction in the tissue boundary. |
| $\overline{\Phi}$ | LEP or GFP fraction. |
| $\overline{\phi_{\text{b}}}$ | LEP or GFP fraction in the tissue boundary. |
| d | Tissue diameter in $\mu\text{m}$. |
| i | Structural macrostates based on $\phi_{\text{b}}$. |
| $p_i$ | Probability of $i^{\text{th}}$ macrostate. |
| $\Omega_i$ | Degeneracy of $i^{\text{th}}$ macrostate. |
| $E_i$ | Mechanical energy of $i^{\text{th}}$ macrostate: |
| $\Delta$E | Mechanical Potential: The derivative of potential energy with $\phi_{\text{b}}$ |
| $\beta$ | Scaled mechanical potential |
| $\gamma_{\text{ll}}$ | Interfacial tension at LEP-LEP interfaces |
| $\gamma_{\text{lm}}$ | Interfacial tension at LEP-MEP interfaces |
| $\gamma_{\text{mm}}$ | Interfacial tension at MEP-MEP interfaces |
| $\gamma_{\text{lx}}$ | Interfacial tension at LEP-ECM interfaces |
| $\gamma_{\text{mx}}$ | Interfacial tension at LEP-ECM interfaces |
| $D_{\text{eff}}$ | Effective diffusion coefficient of cell nuclei |

Table 2: **Specimen information for HMEC.**

| SpecID | Age | Experiment |
|---|---|---|
| 240L | 19 y | Organoid reconstitution, cortical tension, cell-cell and cell-ECM contact angles, nuclear tracking |
| 122L | 66 y | Organoid reconstitution, cell-ECM contact angles |
| 237 | 66 y | Organoid reconstitution, cell-ECM contact angles |
| 029 | 68 y | Organoid reconstitution, cell-ECM contact angles |
| 059L | 23 y | Organoid reconstitution, cell-ECM contact angles |

Table 3: **List of antibodies used.**

| Target | Host | Type | Manufacturer | Product ID | Application |
|---|---|---|---|---|---|
| Keratin-14 | Chicken | Polyclonal | Biolegend | 906001 | IF |
| Keratin-14 | Rabbit | Polyclonal | Biolegend | 905301 | IF |
| Keratin-19 | Rabbit | Monoclonal | ThermoFisher | MA5-31977 | IF |
| Keratin-19 | Mouse | Monoclonal | Biolegend | 628502 | IF |
| Pacific Blue/Ep-CAM | Mouse | Monoclonal | Biolegend | 324218 | FACS |
| APC-Fire 750/CD49f | Rat | Monoclonal | Biolegend | 313632 | FACS |
| APC-Fire 750/CD271 | Mouse | Monoclonal | Biolegend | 345116 | FACS |
| Talin 1 | Rabbit | Monoclonal | Cell Signaling | 4021 | Western blot |
| p120 catenin | Mouse | Monoclonal | BD | 610133 | Western blot |
| $\alpha$-tubulin | Mouse | Monoclonal | Millipore Sigma | T6199 | Western blot |

Table 4: **Morphological parameters recorded for organoid images.**

| Name | Measures | Description |
|---|---|---|
| GFP fraction | composition | Proportion of GFP pixels. |
| Area | shape | Total area in $\mu\mathrm{m}^2$. |
| Circularity | | Roundness, defined as $\frac{4\pi\cdot\text{Area}}{\text{Perimeter}^2}$. |
| Diameter | | Effective diameter of the organoid, $2\sqrt{\text{Area}/\pi}$. |
| Intermixing | cohesion | Number of pixels adjacent to a pixel of the opposite cell type, divided by 2·Area. Higher values mean less cohesion/segregation. |
| Blob Size | | The mean feature size: the reciprocal of the mean fast Fourier transform frequency of the mask with two cell types -1, 1, and 0 elsewhere. |
| Solidity | | Proportion of GFP pixels in the smallest convex enclosing polygon of GFP. Higher values mean more cohesion of GFP. |
| Intercentroid | linear sorting | Distance between the centroids of the cell types, divided by Diameter. Larger values mean more separation. |
| Edge occupancy<br>Outer occupancy<br>Inner occupancy | radial sorting | Proportion of GFP in Edge (directly adjacent to ECM) or Outer/Inner (lower/upper third of pixels ranked by distance to ECM). |
| Correctness | | Proportion of pixels that match the segmentation generated by moving all MEP pixels as close to the ECM as possible. |
